# Contrasting relationships between evolved consumer diversity and productivity for essential vs. substitutable resources

**DOI:** 10.64898/2026.10.08.757349

**Authors:** Aswin Gopakumar, Sebastian Diehl, Åke Brännström

## Abstract

Any mechanistic biodiversity-ecosystem functioning theory must explain how biodiversity in natural communities arises, which requires an evolutionary metacommunity perspective. Using a model of adaptive evolution of competitive metacommunities in landscapes with spatially heterogeneous supply of two resources, we studied the evolution of resource-uptake specialization as a function of resource type (substitutable/essential) and trade-off shape between resource-uptake affinities (generalist-/specialist-favouring). We identified ‘critical essentiality’ as a key phenomenon separating parameter regions with fundamentally different biodiversity-ecosystem function relationships. Specifically, while greater spatial heterogeneity in resource supply increases biodiversity, this comes at the cost of reduced productivity when resources are essential. Moreover, when resources are essential, both generalist- and specialist-favoring trade-offs select for phenotypes that prevent efficient resource partitioning, leading to low resource use efficiency and consumer biomass – despite often high species richness. In contrast, when resources are substitutable, a specialist-favoring affinity trade-off selects for phenotypes that partition resources efficiently, promoting a metacommunity of only two species characterized by high functional diversity, resource use efficiency and biomass. Overall, our analyses thus suggest that conditions promoting the evolution of diverse consumer communities often do not engender higher functional richness, consumer biomass, or resource use efficiency.

## 1 Introduction

A fundamental question in ecology is how the productivity of ecosystems relates to species diversity. A common perception is that higher levels of consumer diversity should result in higher resource use efficiency, i.e., in stronger depletion of the supplied resources and a concomitant increase in consumer production and biomass (Ptacnik et al. 2008; Plas 2019; Hodapp et al. 2019; Wang et al. 2024). This perception has empirical support from experimental studies with randomly assembled communities, in particular grassland communities in which diversity treatments were typically maintained through weeding (Balvanera et al. 2006; Cardinale et al. 2011; Tilman et al. 2014). Yet, it has also been noticed that diversity-productivity relationships can differ between experiments and observational studies, with the latter often showing unimodal or even negative relationships between productivity and diversity (Mittelbach et al. 2001; Thompson et al. 2005; Grace et al. 2007; Jiang et al. 2009; Fraser et al. 2015; Plas 2019). Moreover, biodiversity manipulations that were left open to colonization by the surrounding species pool have frequently developed negative diversity-productivity relationships (Pfisterer et al. 2004; Petermann et al. 2010; Rychtecká et al. 2014).

The apparent discrepancy between the biodiversity-productivity patterns found in controlled experiments versus observational studies and experiments that were open to colonization-extinction processes suggests that both abiotic spatial heterogeneity and the regional species pool contribute to shaping the relationship between diversity, productivity and resource use efficiency in natural communities (Loreau et al. 2001; Ptacnik et al. 2010; Hodapp et al. 2016). To address this issue, mathematical models have been developed that explore the interplay of spatial variation in habitat quality or resource supply with regional metacommunity dynamics, and their combined impact on diversity-productivity relationships. Such metacommunity-based theoretical approaches have indeed confirmed the possibility of various shapes of the diversity-productivity relationship at both local and regional scales (Kondoh 2001; Mouquet et al. 2003; Gross et al. 2007; Thompson et al. 2016; Hodapp et al. 2016; Thompson et al. 2020). While these studies make important steps forward towards reconciling seemingly conflicting empirical results, they all focus on purely ecological dynamics, typically using arbitrary or randomly drawn species pools and trait combinations. Yet, a truly synthetic diversity-productivity theory needs to explain how biodiversity is generated and maintained in the first place, which requires an evolutionary perspective (Matthews et al. 2011; Mayfield et al. 2023). Such a perspective is likely to modify previous conclusions, because eco-evolutionary models commonly predict that the species richness at an evolutionary endpoint is considerably lower than the number of species that could coexist ecologically (Koffel et al. 2016; Edwards et al. 2018).

As a first step towards the development of an evolutionary productivity-diversity theory, we used an adaptive dynamics approach of the evolution of competitive metacommunities in landscapes with spatially heterogeneous resource supply (Wickman et al. 2017) to explore how the species richness of adaptively evolved metacommunities is related to functional diversity, resource use efficiency, and community biomass. In a previous study, Wickman et al. (2019) introduced the model to study how the evolved species richness of communities of consumers of two potentially limiting resources depended on (i) the shape of the trade-off between the evolving traits, i.e., the consumers’ uptake affinities for the two resources (from generalist-favoring to specialist-favoring) and (ii) the degree of resource substitutability (from substitutable to essential). They found that the conditions promoting evolutionarily stable coexistence in homogeneous environments, namely substitutable resources and specialist-favoring trade-offs, gave rise to among the least diverse metacommunities in heterogeneous landscapes, where the most diverse communities instead evolved for clearly essential resources and intermediate trade-offs. Their analysis stopped, however, at the evolved diversity, and did not explore the relationships between diversity and productivity that emerged.

Here, we expand on the analysis by Wickman et al. (2019) and ask the following questions: (1) How do species richness, biomass, and resource use efficiency of evolved metacommunities of resource competitors depend on the shape of the affinity trade-off and the degree of resource substitutability? (2) Do conditions that promote the evolution of diverse communities also engender higher community biomass and resource use efficiency? (3) Given that ecosystem functions such as productivity and resource use efficiency are not causally linked to species richness per se but to the traits of the species in a community (Hillebrand et al. 2009) – does evolved species richness correlate with the functional richness of evolved communities? (4) Does adaptive evolution lead to transgressive overyielding, i.e., does it increase community biomass and resource use efficiency compared to the best-performing monoculture? (5) Given that spatial heterogeneity in resource supply ratios can cause local imbalances between resource supply and demand (Hodapp et al. 2016; Leibold et al. 2017) – how does spatial heterogeneity in resource supply affect the biomass and resource use efficiency of evolved communities, and how does this depend on the shape of the affinity trade-off and the degree of resource substitutability?

## 2 Methods

### Demographic dynamics

We use the consumer-resource model developed by Wickman et al. (2019), in which *N* consumer species with densities *A_i_* (*i* = 1*, …, N*) feed on two resources with densities *R*_1_ and *R*_2_ in a spatially heterogeneous, rectangular landscape with periodic boundary conditions. Consumers and resources disperse through diffusion (the macroscopic limit of independent Brownian motion), while spatial heterogeneity arises from local differences in resource supply, resource consumption, and consumer growth. Local system dynamics are described by the following partial differential equations:

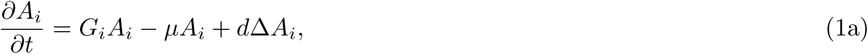

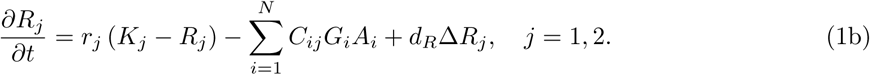

State variables, functions, and parameters are defined in Table 1 and briefly described below. The terms in the consumer rate of change are biomass gains from resource consumption at rate *G_i_A_i_*, losses from mortality at rate *µA_i_*, and net dispersal at rate *d*. The terms in the resource rate of change are gains from resource renewal, losses from consumption, and net dispersal – where *r_j_* is the renewal rate, *K_j_* the local supply density, *C_ij_* the fractional contribution to the diet and growth of consumer *i*, and *d_R_* the diffusion rate of resource *j*. Consumer growth rates *G_i_* and diet contributions *C_ij_* are functions of other model components, as we specify next.

**Table 1:**
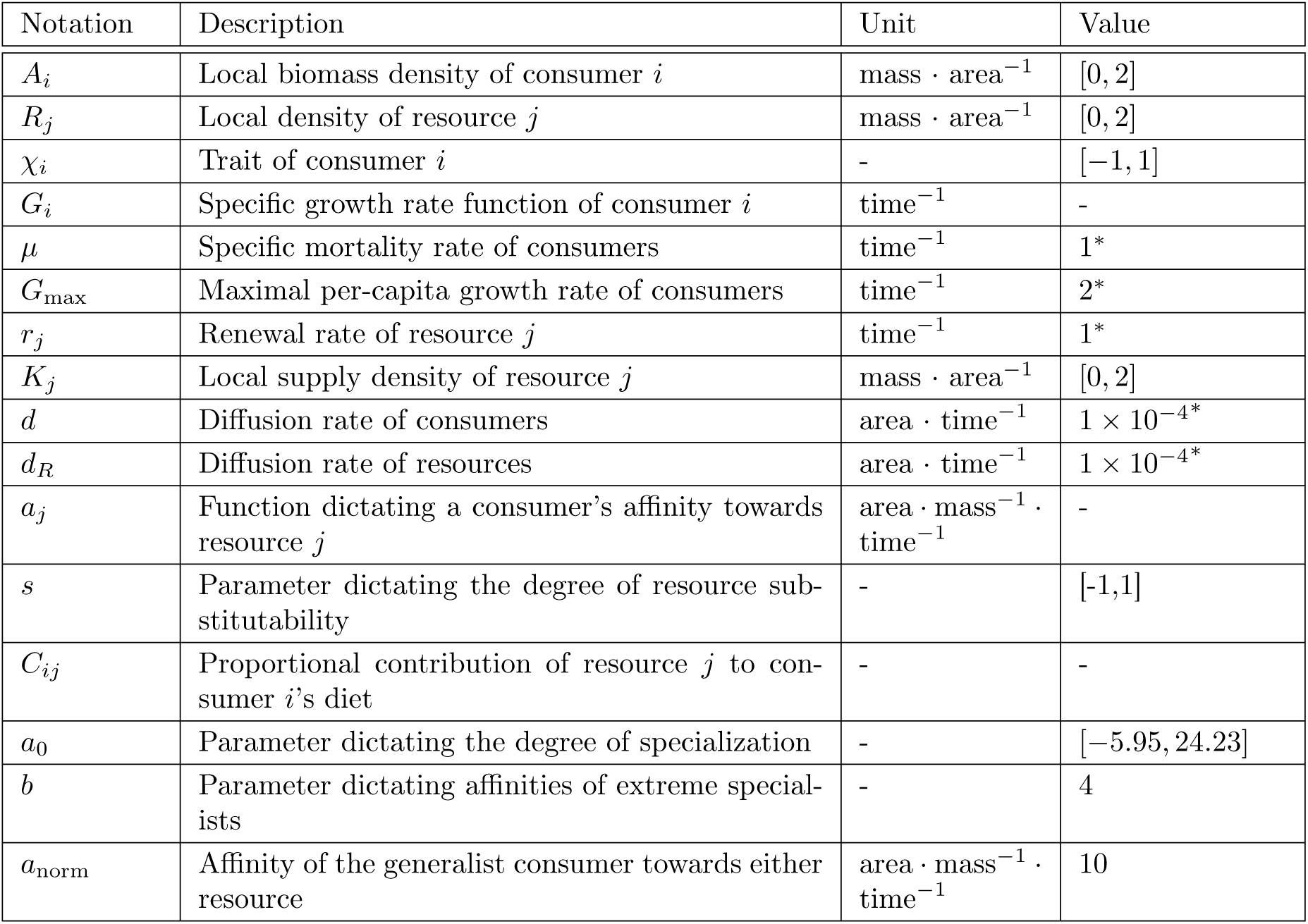
Definitions of state variables, functions, and parameters. The last column lists the possible minimum and maximum values of state variables and the values of parameters that were either held constant across all simulations or varied within the ranges in brackets in diflerent simulations, respectively. Values with asterisks are derived from the non-dimensionalized parameters used for the simulations. More information on the non-dimensionalization and parametrization of the non-dimensionalized system is provided in the appendix.

### Resource consumption

Each consumer species *i* is characterized by a trait value *χ_i_*that determines its uptake affinities *a*_1_(*χ_i_*) and *a*_2_(*χ_i_*) for the two resources. A consumer’s per-capita growth rate, *G_i_*, is a function of these affinities, the local resource densities, and a tunable parameter *s* that allows resources to range continuously from substitutable to essential (León et al. 1975; Tilman 1980):

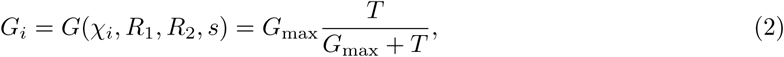

where

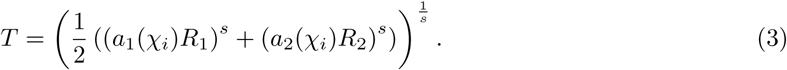

Here, the products *a*_1_(*χ_i_*)*R*_1_ and *a*_2_(*χ_i_*)*R*_2_ are the ‘effective resource availabilities’ (Wickman et al. 2019), and the value of *s*, *−∞ < s ≤* 1, determines to what extent resources are substitutable (Schreiber et al. 2003). Thus, when *s* = 1, resources are perfectly substitutable, and the per-capita growth rate assumes the form of a standard Holling type II functional response with handling time *h* = 1*/*(2 *G*_max_):

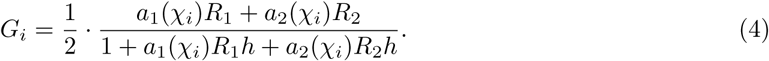

On the other extreme, when *s → −∞*, resources are strictly essential, and the per-capita growth rate assumes the form of a Monod-type minimum function

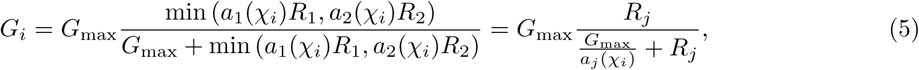

with half-saturation constant *G*_max_*/a_j_*(*χ_i_*), where *a_j_*(*χ_i_*) is the consumer’s affinity for its currently growth limiting resource *j*.

We set the conversion efficiency of consumed resources equal to 1, which is without loss of generality provided it is identical across consumer species and both resources, as it can otherwise be absorbed into the scaling of biomass and resource densities. We can further impose the mass-balance condition that the proportional contributions of the two resources to a consumer’s consumption sum to unity, *C_i_*_1_ + *C_i_*_2_ = 1, so that one unit of consumer growth removes one unit of resource biomass in total, irrespective of the degree of substitutability. This condition underlies the equilibrium relationship between consumer biomass and residual resource density derived in the appendix.

How the consumer’s consumption is divided between the two resources differs at opposite ends of the substitutability spectrum. When resources are perfectly substitutable, the consumers’ growth rate depends on the sum of the effective resource availabilities, and each resource is consumed in proportion to its share of that sum, *C_ij_*= (*a_j_*(*χ_i_*)*R_j_*)*/*(*a*_1_(*χ_i_*)*R*_1_ + *a*_2_(*χ_i_*)*R*_2_). When resources are strictly essential, growth is set by the limiting resource alone, and the two resources are instead consumed in the fixed ratio required to build new biomass, which under our symmetry assumptions is *C_i_*_1_ = *C_i_*_2_ = 0.5.

For intermediate resource types, we interpolate continuously between these two limits,

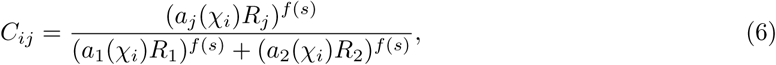

where *f* (*s*) = exp(*s −* 1) is an increasing function that satisfies *f* (*−∞*) = 0 and *f* (1) = 1. This functional form for intermediate substitutability is phenomenological rather than mechanistically derived, but these choices ensure that when resources are not strictly essential, a consumer’s resource consumption is skewed towards the one with higher effective availability (Wickman et al. 2019).

### Consumer affinity trade-off

The affinities *a*_1_ and *a*_2_ determine the ability of a consumer to acquire the two resources. We assume that consumers can only increase their affinity for one resource at the expense of decreasing their affinity for the other one, following a fixed trade-off curve that may favour either resource generalists or resource specialists (see fig. 2). Any consumer’s resource affinities are determined by its trait value *χ* as

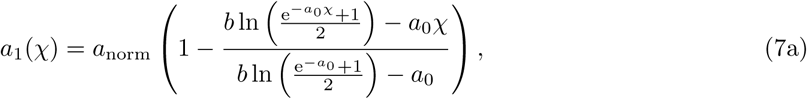

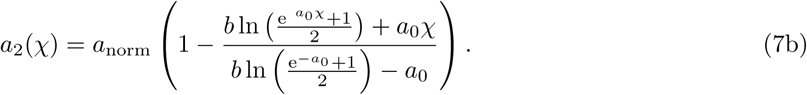

**Figure 1:**
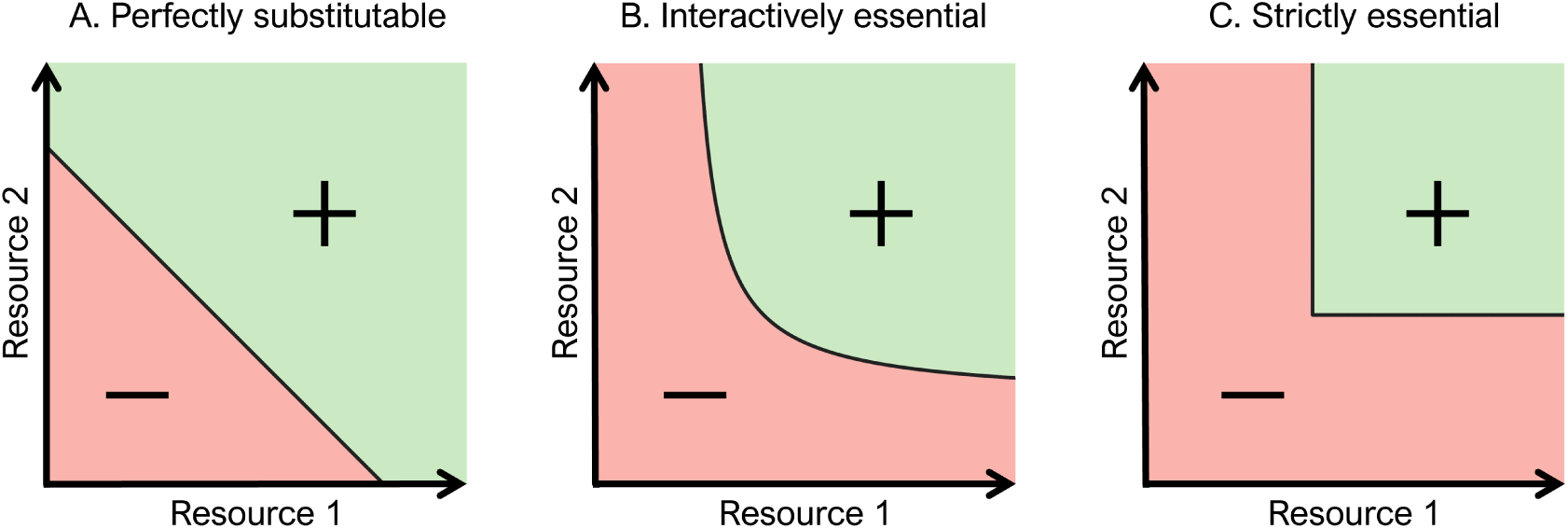
The spectrum of resource substitutability illustrated with zero net-growth isoclines of the resource generalist (consumer with a_1_ = a_2_ = a_norm_, black lines) for: (A) perfectly substitutable resources (s = 1), (B) interactively essential resources (s = −1), and (C) strictly essential resources (s = −∞). Green and red regions indicate combinations of resource densities for which the resource generalist has positive and negative growth, respectively.

**Figure 2:**
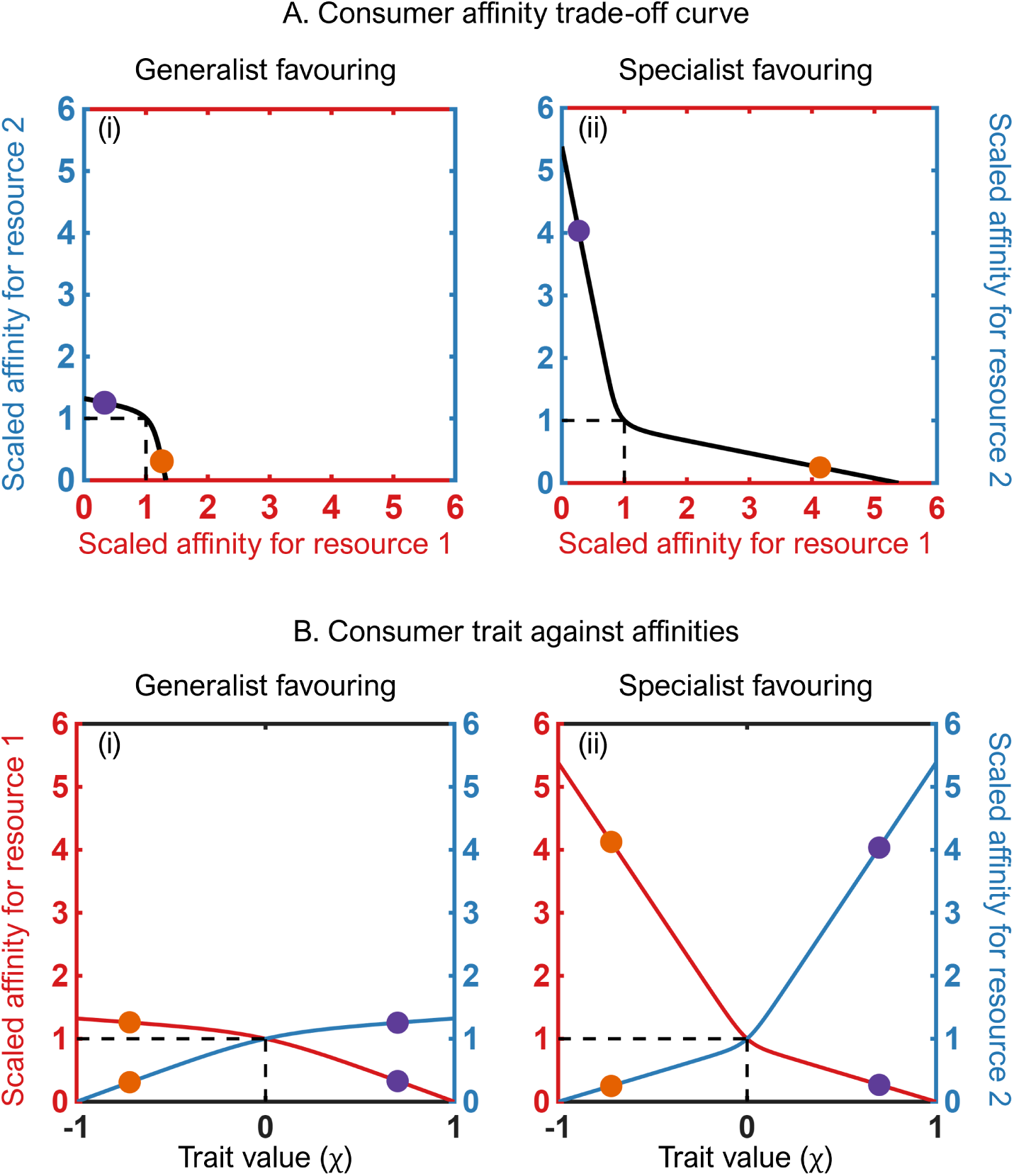
Examples of affinity trade-offs and resulting consumer trait combinations. (A) Scaled consumer affinity trade-off curves ((â_1_, â_2_) = (a_1_/a_norm_, a_2_/a_norm_)) illustrating (i) the most generalist-favoring trade-off (a_0_ = −5.95, b = 4) and (ii) the most specialist-favoring trade-off (a_0_ = 24.23, b = 4) used in numerical simulations. (B) Corresponding functions relating scaled resource affinities â_1_and â_2_ to the consumer trait value, χ, for (i) the generalist-favoring trade-off and (ii) the specialist-favoring trade-off from panel A. Red and blue curves in panels B(i) and B(ii) represent the affinities for resource 1 and resource 2 at a given trait value χ, respectively. Orange and purple dots correspond to consumers with trait χ = −0.7 and χ = 0.7, respectively. Broken lines mark the scaled affinities of a perfect resource generalist ((â_1_, â_2_) = (1, 1)), corresponding to a trait value χ = 0.

Here, the parameter *b* constrains the maximum affinities of extreme specialists (*χ* = 1 and *χ* = *−*1, respectively), while the parameter *a*_0_ can be tuned to change the trade-off continuously from generalist-favoring to specialist-favoring. The strength of the trade-off is measured using the curvature of the consumer affinity trade-off curve near the resource generalist (panels (i) and (ii) of fig. 2A). Note that the trade-off curve always passes through the point (*a*_1_*, a*_2_) = (*a*_norm_*, a*_norm_) to ensure that a generalist consumer is always comparable between the scenarios considered.

### Evolutionary dynamics

Evolutionary dynamics were modeled using adaptive dynamics techniques, which center on the concept of invasion fitness, defined as the growth rate of an initially rare mutant subpopulation invading a community of one or more resident populations (Metz et al. 1996; Geritz et al. 1998; Dieckmann et al. 1996; see also Brännström et al. 2013). Mutant and resident(s) are characterized by their respective values of the affinity-determining trait *χ_i_*. Mutations of the trait are assumed to be rare, so that the resident community can be assumed to be at demographic equilibrium when a mutation occurs. A mutant subpopulation can invade if its trait value conveys a selective advantage, i.e., its invasion fitness is positive. Assuming that mutational steps are small, we describe directional selection on the trait with a differential equation for each consumer,

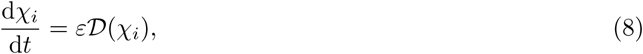

where *D*(*χ_i_*) denotes the selection gradient of consumer *i* in the current resident environment, and *ε* = 10*^−^*^6^ determines the time scale separation between the ecological and evolutionary dynamics. The selection gradient is the slope of the invasion fitness around the trait value *χ_i_* of consumer *i*, which, in a spatially heterogeneous environment, can be determined using the method introduced by Wickman et al. (2017).

In this study, we initiate the evolutionary community assembly process with a single, near-perfect generalist consumer (Fig. 3A) and solve the equations for the ecological dynamics (eqn. 1) until the system has reached demographic equilibrium. We then simultaneously solve the equations for the ecological dynamics and the trait dynamics (eqn. 8) until the selection gradient at the resident consumer’s trait value becomes zero (black dot in panel (i) of fig. 3B). The system is then said to have reached an evolutionarily singular state. To determine whether further mutant invasions are possible, the invasion fitness is calculated for all possible mutant trait values. If there are mutant trait values with positive invasion fitness, we let the mutant with the highest invasion fitness invade the system (red dot in panel (i) and green dot in panel (ii) of fig. 3B). This cycle of mutant invasion followed by evolutionary trait adaptation is repeated until there is no trait space left with positive invasion fitness (panel (iii) of fig. 3B), and an evolutionarily steady state has been reached where the community is closed to further invasions. Extensive numerical explorations have shown that this evolutionary endpoint is unique in all cases considered in our study (Wickman et al. 2019).

**Figure 3:**
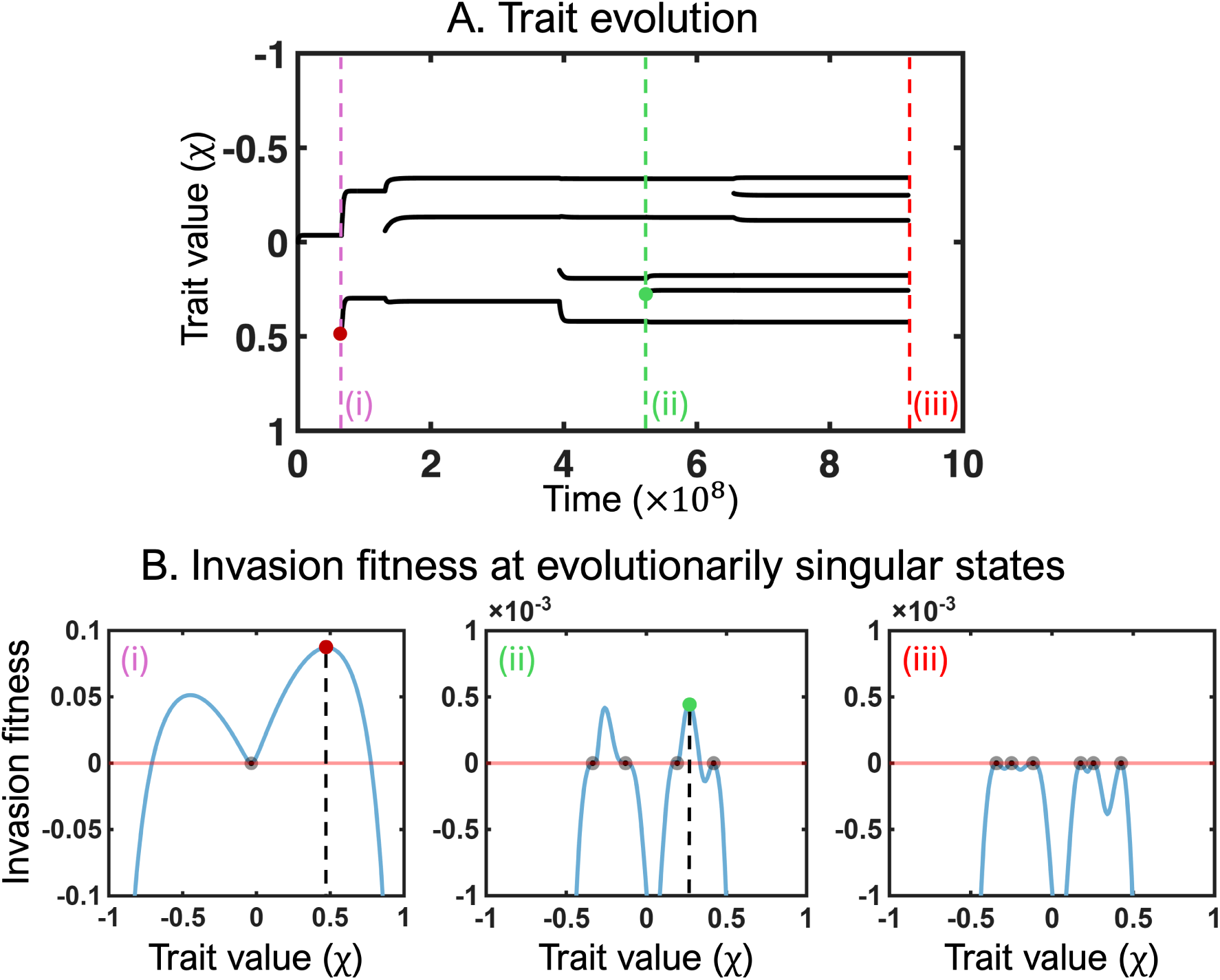
Graphical representation of the adaptive dynamics methodology used in modeling evolutionary dynamics. (A) Time course of the interplay of continuous trait evolution and occasional mutant invasion leading to an evolutionarily steady state with 6 consumer species. (B) Invasion fitness of residents (black dots) and successfully invading mutants (colored dots) as a function of their trait values at the three events highlighted by broken lines in panel A: (i) first speciation, (ii) fourth speciation, and (iii) evolutionarily steady state. Parameter values used in the simulation are: generalist favoring trade-off with a_0_ = −0.17, and interactively essential resource type with s = −1.

### Experimental design and analytical procedures

We explored the combined influences of resource substitutability and the curvature of the affinity trade-off on the biomass, diversity and trait distribution of evolved consumer communities in a full factorial design, in which we varied the degree of resource substitutability in 18 steps from perfectly substitutable (*s* = 1, Fig. 1A) to interactively essential (*s* = *−*1, Fig. 1B), and the shape of the consumer affinity trade-off curve in 54 steps from strongly generalist-favoring (*a*_0_ *≈ −*5.95) to strongly specialist-favoring (*a*_0_ *≈* 24.23). The steps of *a*_0_ were chosen so that the curvature of the trade-off curve, evaluated at [*a*_1_*, a*_2_] = [1, 1], ranged from strongly generalist-favoring (Fig. 2A(i)) to strongly specialist-favoring (Fig. 2A(ii)). More extreme trade-offs seemed biologically unreasonable, while simulations with strictly essential resources proved to be computationally prohibitive, at least for the relatively low diffusion rates that we used in this study. Additional simulations performed at higher resource essentiality combined with higher diffusion rates did not, however, reveal any qualitatively different results and are therefore not shown.

We used Perlin noise (Perlin 1985) to create periodic, spatially heterogeneous resource supply landscapes on a 32×32 grid, on which the resource supplies *K*_1_ and *K*_2_ were varied in a spatially autocorrelated manner. All simulations were run on four replicate realizations of each of three types of landscapes that differed in the degree of cross-correlation between the resource supply densities *K*_1_ and *K*_2_ – perfectly anti-correlated (*ρ* = *−*1), uncorrelated (*ρ* = 0) and strongly correlated (*ρ* = 0.95) – yielding a total of 11, 664 evolutionary community assembly simulations. To more precisely describe the trait distributions of evolved communities across randomly generated landscapes, we performed additional simulations combining three levels of resource substitutability (*s ∈ {−*1, 0, 1}) with four widely spaced shapes of the affinity trade-off, now using 160 replicate landscape realizations of each of the three types of cross-correlation between *K*_1_ and *K*_2_. To be used in any of the numerical experiments, a landscape had to fulfil three criteria: (1) the average supply of both resources was equal, i.e. ∫ *K*_1_ d*x* = ∫ *K* _2_ d*x* = 1; (2) spatial autocorrelation of both *K*_1_ and *K*_2_, measured as Moran’s *I*, ranged from 0.175 to 0.2; and (3) the coefficient of variation (CV) in both *K*_1_ and *K*_2_ ranged from 0.3 to 0.325.

Each simulation was initiated with a single, near-perfect generalist consumer and run to evolutionarily steady state (see example in fig. 3 and 4B). During community assembly, the successful invasion of a mutant can sometimes cause the extinction of a resident. We considered a species extinct when its biomass had declined below 10*^−^*^6^ everywhere on the landscape. Once a simulation had reached the evolutionarily steady state, we recorded the number of consumer species (i.e., species richness), their traits and their biomass, as well as the abundances of the two resources.

**Figure 4:**
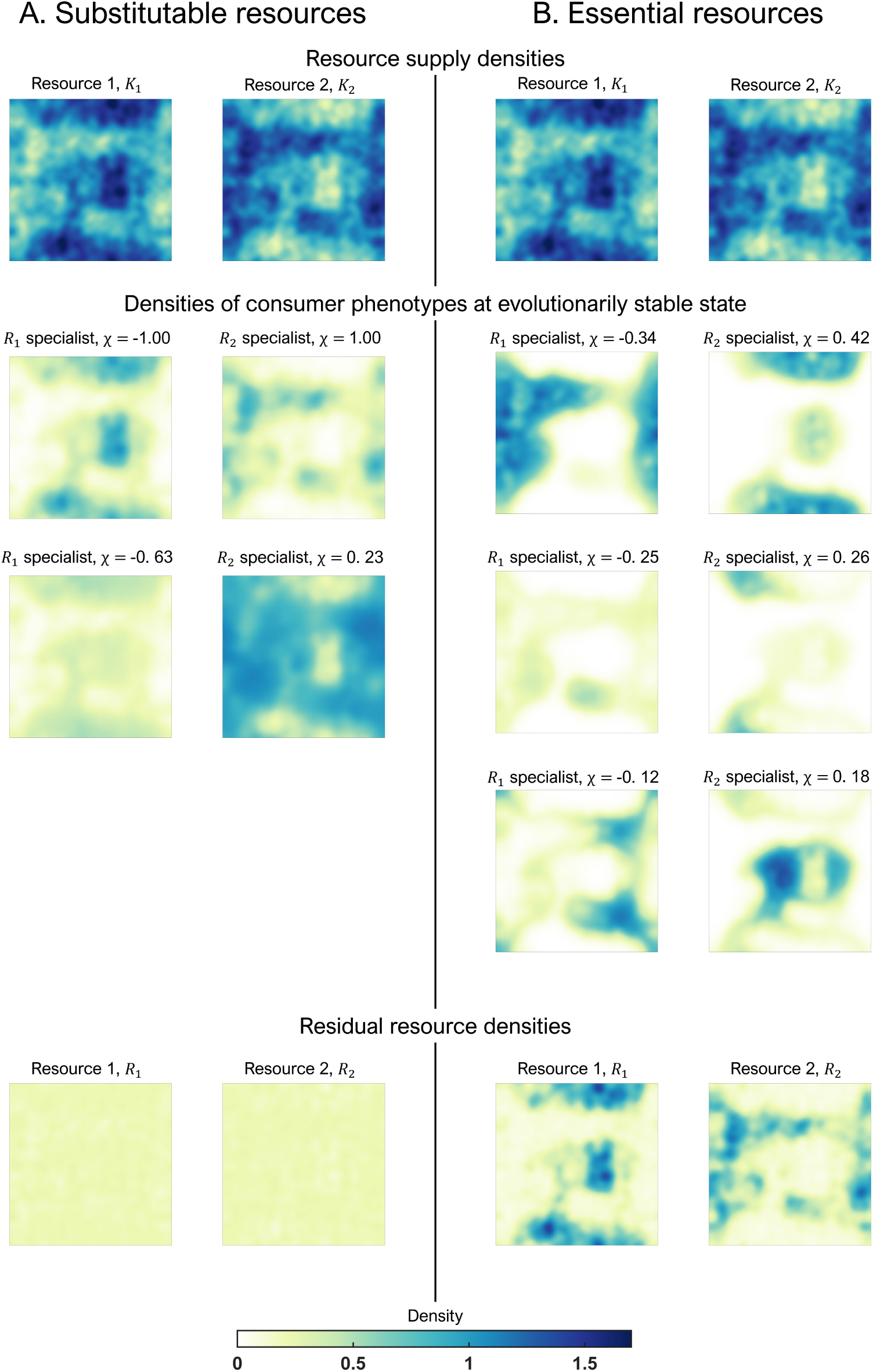
Two examples of the trait values and spatial distributions of two diflerent consumer communities and their resources at an evolutionarily steady state when resources are (A) substitutable and (B) essential. Top panels: Spatial distribution of the resource supply densities K_1_and K_2_ in the (perfectly anti-correlated) resource supply landscape on which the two communities evolved. Middle panels: Trait values (χ) and spatial density distribution of consumer species at the evolutionary endpoints. Consumers are ranked in descending order of specialization, with the strongest resource 1 and resource 2 specialists at the top. Bottom panels: Corresponding resource densities (R_1_ and R_2_) at the evolutionary endpoints. Simulations assumed a mildly generalist favoring trade-off (a_0_= −0.17) and either perfectly substitutable resources (s = 1, left columns of panels) or interactively essential resources (s = −1, right columns of panels), corresponding to Roman numerals ii and x in panel A of Fig. 5, respectively. Simulations were run on a 32 × 32 grid and upscaled to a 128 × 128 grid with a periodic interpolation program for visualization.

We subsequently assessed functional richness and functional divergence of the evolved community. Functional richness was assessed as the volume of the occupied trait space, expressed as the Euclidean distance between the affinities of the species with the most extreme trait values

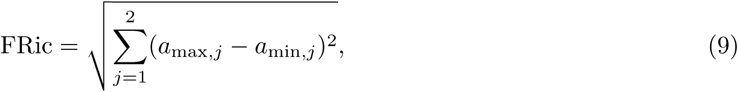

where *a*_max_*_,j_* and *a*_min_*_,j_* are, respectively, the highest and lowest affinities towards resource *j* observed in the evolved community. The above measure of functional richness does not take into account how well the trait space between the extremes is filled by the community and can give biased results if the species with the most extreme trait values contribute a minor proportion to total community biomass. We therefore also determined functional divergence as Rao’s entropy (Rao 1982; Botta-Dukát 2005)

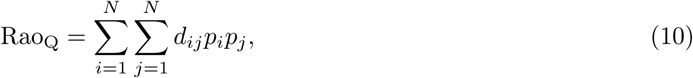

where *d_ij_* is the Euclidean distance between consumer *i* and *j* in the trait space, and *p_i_* and *p_j_* are the fractional biomass of consumer *i* and *j* in the community, respectively.

For each evolved species *i*, we furthermore calculated its landscape-wide ‘trait-resource alignment’ (TRA*_i_*), which is a measure of the alignment of the species’ spatial distribution with the supply of the resource towards which it has the higher affinity. Following Wickman et al. (2019, Appendix S1), we calculated the trait-resource alignment of species *i* on a given landscape as

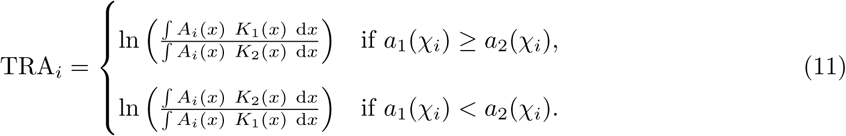

where *A_i_*(*x*) is the biomass of species *i* at location *x*, and *K*_1_(*x*) and *K*_2_(*x*) are the supply densities of the two resources at that location. The value of TRA*_i_* is positive, if a resource 1 specialist has a higher biomass in locations where the supply of resource 1 is high compared to locations where the supply of resource 2 high, and negative otherwise. The community-wide overall trait-resource alignment, which we introduce here, is then given by the biomass-weighted, average trait-resource alignment of all species in the community i.e.,

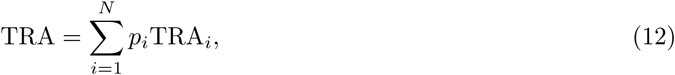

where *N* is the species richness of the community and *p_i_* is the proportional biomass of species *i*.

We also determined the landscape-wide resource use efficiency of the evolved communities, which we defined as

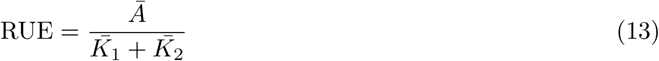

where the numerator and denominator represent total consumer biomass 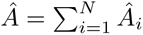 and total resource supply, respectively, averaged across the landscape. Since 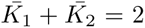 in all simulations, and the sum of total consumer biomass and the residual resource densities 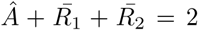, we subsequently use consumer biomass synonymously with resource use efficiency.

We finally investigated how an important ecosystem function, i.e. the landscape-averaged biomass production of the evolved consumer community 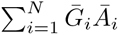 (a) compares to the production of the corresponding most productive monoculture, and (b) is affected by the spatial heterogeneity of the resource supply landscape. Since, at demographic equilibrium, the landscape-averaged production of each consumer species must equal the average biomass multiplied by the constant per-capita mortality rate 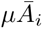, we used average total consumer biomass as a proxy that is directly proportional to total community production. For comparison with the most productive monoculture in a given resource supply landscape, we searched the trait space to identify the monomorphic consumer that yielded the highest biomass at demographic equilibrium. For each combination of given resource substitutability and affinity trade-off, we then quantified the relative biomass difference between the evolved community and the most productive monoculture as

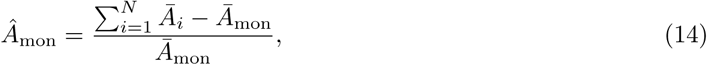

where *N* is the species richness in the community and *Ā*mon is the total consumer biomass of the most productive monoculture.

To assess the consequences of spatial heterogeneity in resource supply for biomass production, we repeated the evolutionary community assembly experiments in a spatially homogeneous environment with an overall resource supply that was identical to that in the heterogeneous landscapes (i.e., *K*_1_ = *K*_2_ = 1). For each combination of a given resource substitutability and affinity trade-off, we then quantified the effect of spatial heterogeneity as

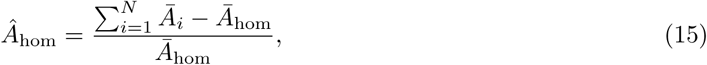

where *Ā*hom is the total consumer biomass of the evolved community in the homogeneous environment.

## 3 Results

The degree of resource substitutability and the shape of the affinity trade-off, as well as their interaction, have clear and systematic effects on the diversity, biomass, and resource use efficiency of evolved metacommunities. These effects are observed in all resource supply landscapes, but are typically most strongly expressed in landscapes in which the local supply densities of the two resources are negatively cross-correlated. In the following, we therefore use negatively cross-correlated resource supply landscapes to illustrate the main results. Corresponding figures for landscapes with uncorrelated or positively correlated resource supply densities are shown in the appendix, and trends along the cross-correlation gradient will be described at the end of the results section.

### Critical essentiality: a sharp divide, separating evolutionary responses to the supply with substitutable versus essential resources

A striking feature of our numerical results is a qualitative transition, at a single point along the continuous gradient in resource substitutability, between two contrasting regimes of evolutionary outcomes, one for substitutable-like and one for essential-like resources. This transition corresponds with a pronounced minimum in the diversity of evolved consumers (Figs. 5-7; also see the appendix). The boundary, called ‘critical essentiality’ by Wickman et al. (2019), arises because, on either side of the boundary, consumers show contrasting trait-resource alignment with the supply of the resource for which they have the higher affinity. Thus, when resources are substitutable, a resource 1 specialist is locally most competitive in landscape patches where the supply is biased towards resource 1, i.e. *^K^*^1^*/K*2 *>* 1 (Fig. ^4^A). In contrast, when resources are essential, a resource 1 specialist is locally most competitive in landscape patches where the supply is biased against resource 1, i.e. *^K^*^1^*/K*2 *<* 1 (Fig. ^4^B). This latter apparent ‘misalignment’ arises because, in landscape patches where *^K^*^1^*/K*2 *<* 1, the relatively higher supply of resource 2 releases the resource 1 specialist from limitation by resource 2, thus enabling it to depress resource 1 to levels where resource 2 specialists cannot persist. The reverse reasoning applies to resource 2 specialists, which are most competitive for substitutable resources where the supply is biased towards resource 2 ( *^K^*^2^*/K*1 *>* 1; Fig. ^4^A), and most competitive for essential resources where the supply is biased against resource 2 (*^K^*^2^*/K*1 *<* 1; Fig. ^4^B). As described in the following sections, this qualitative shift in trait-resource alignment at the transition from substitutable to essential resources has profound consequences for species richness, trait distribution and biomass of the evolved consumer communities, yielding contrasting patterns in all response variables above vs. below the critical essentiality boundary (Figs. 5-7).

**Figure 5:**
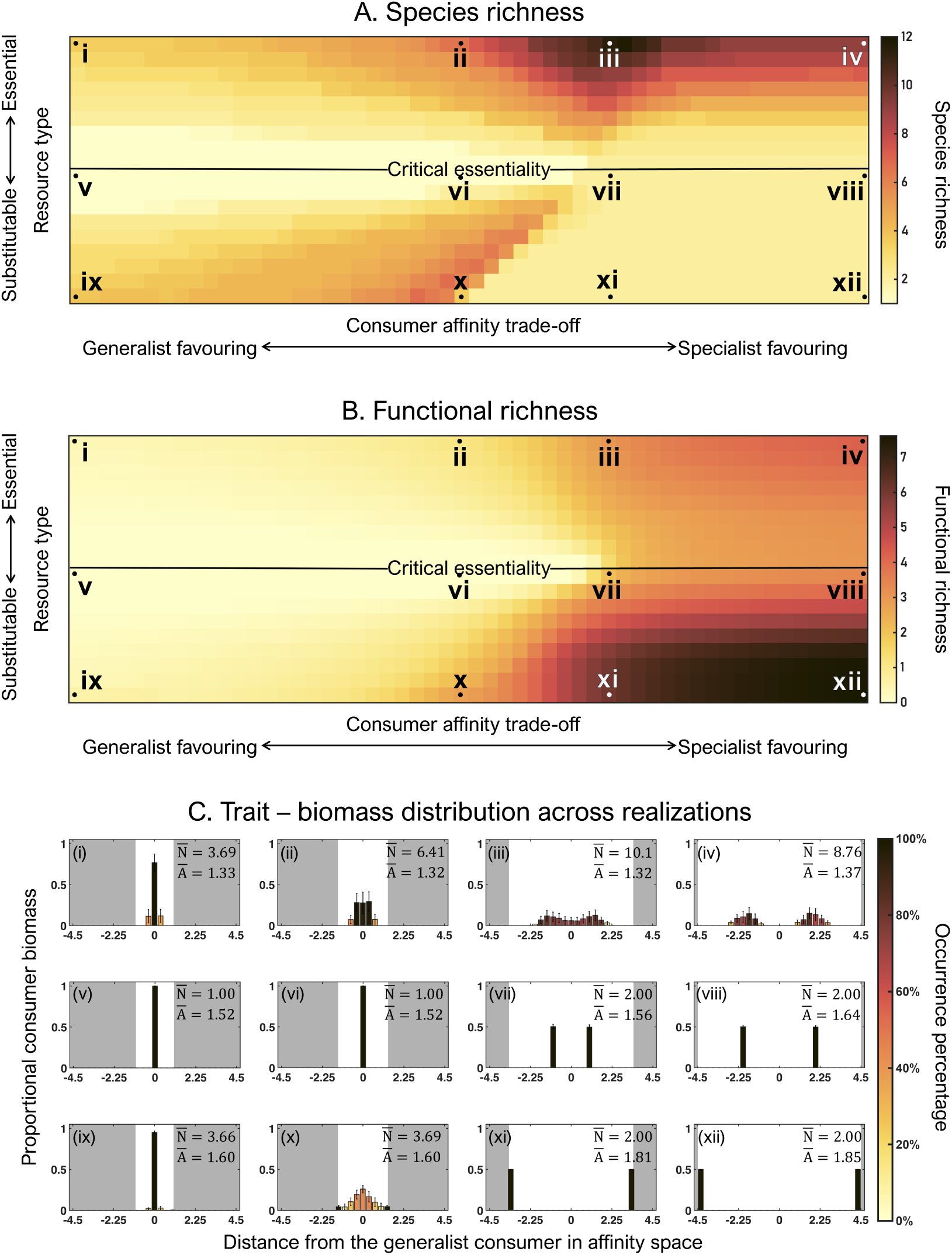
Evolved (A) species richness, and (B) functional richness and (C) trait distribution in landscapes with perfectly anti-correlated (ρ = −1) supplies of resources 1 and 2 and for diflerent combinations of the strength of the affinity trade-off (ranging from strongly generalist- to strongly specialist-favoring, see fig. 2A) and the degree of resource substitutability (ranging from perfectly substitutable to interactively essential, see Fig. 1A and B). Panels A and B show mean values of four randomly generated resource supply landscapes, with the central horizontal black lines indicating the critical resource essentiality. Histograms in panel C show the distribution of consumer biomass in affinity trait space of evolved metacom munities across 160 randomly generated landscapes for the 12 combinations of affinity trade-off strength and degree of resource substitutability indicated by Roman numerals in panels A and B. N and Ā indicate the average values of species richness and total consumer biomass across all 160 realizations. Columns represent the average proportional biomass (+1 standard deviation) of consumers that are specialists on resource 1 (negative x-axis values) and resource 2 (positive x-axis values), respectively, where the histograms’ x-axes represent the specialists’ distance in affinity space from the generalist consumer (with affinities a_1_= a _2_ = 1), with higher absolute values indicating stronger specialization, and the bar color indicating the frequency of occurrence of an affinity distance across all 160 realizations. The white space around each histogram indicates the available trait space, where borders to the gray areas mark the maximum affinity values on a given trade-off curve.

### Species rich communities can only evolve when the resources are clearly essential or clearly substitutable

On the continuous gradient in resource substitutability, species richness of evolved consumer communities is always extremely low near the critical essentiality, where resources are neither clearly substitutable nor clearly essential (positions v-viii in Fig. 5A). This is because, at the critical essentiality, a species’ competitiveness is defined by the product *a*_1_(*χ_i_*) *· a*_2_(*χ_i_*) of its resource affinities (Wickman et al. 2019). This product is highest for a perfect generalist when the affinity trade-off favors generalists or mild specialists and for one medium specialist on either resource when the affinity trade-off favors stronger specialists. Near the critical essentiality, evolved communities therefore consist of either a monoculture of the perfect generalist (generalist-favoring trade-off, panels v-vi in Fig. 5C) or a complementary pair of medium specialists (specialist-favoring trade-off, panels vii-viii in Fig. 5C).

When resources are clearly substitutable or clearly essential, species-rich communities become possible (positions i-iv and ix-xii in Fig. 5A), the evolved species richness being highest when the consumer affinity trade-off is neither strongly generalist-nor strongly specialist-favoring (vicinities of positions iii and x in Fig. 5A). When the affinity trade-off favors generalists strongly, the community is dominated by a small number of resource generalists, and more strongly specialized species cannot evolve (panels i and ix in Fig. 5C). Moving from the strongly generalist-towards a more specialist-favoring trade-off, the expanding trait space is initially filled by an increasing number of mild specialists, yielding both an increase in species richness and an increasingly broad, unimodal trait distribution centered on the perfect generalist (panels ii and x in Fig. 5C).

Moving further towards a specialist-favoring trade-off weakens the competitiveness of generalists, and the trait distribution becomes first bimodal and then discontinuous, with a widening gap around the perfect generalist’s trait combination. When resources are substitutable, species richness declines abruptly upon the disappearance of generalists, and community composition shifts to a single pair of strong and complementary specialists (panels xi-xii in Fig. 5C). Since resources are substitutable, these two strong specialists can persist everywhere in the landscape and, together, drive resources to levels that are too low for any other affinity combination to exist. In contrast, when resources are essential, resource monopolization by the same pair of specialists across the entire landscape is not possible, because different supply ratios select locally for different affinity combinations. When resources are essential, species richness therefore declines more gradually upon the disappearance of generalists (panels iii-iv in Fig. 5C).

### Evolved species richness is a poor predictor of evolved functional richness

Functional richness (FRic) and functional divergence (Rao’s entropy) were highly correlated across all 11,664 simulations (*R*^2^ = 0.99). We therefore only show results based on functional richness, with the results for Rao’s entropy being nearly indistinguishable.

Moving from a strongly generalist- to a strongly specialist-favoring trade-off, the possible trait space (indicated by the white trait windows in Fig. 5C) continuously increases. Not surprisingly, functional richness of evolved communities therefore increases monotonically along this gradient. In consequence, functional richness and species richness are poorly correlated, since species richness has a unimodal relationship with the affinity trade-off (compare panels A and B in Fig. 5). In fact, the overall highest functional richness is observed for a species richness of only two species, i.e., when a strongly specialist-favoring trade-off promotes the evolution of a community of two complementary, extreme resource specialists on perfectly substitutable resources (community xii in Fig. 5A-C). In contrast, the more diverse communities of consumers of essential resources cannot sustain extreme specialists on a single resource, and therefore cannot reach similarly high functional diversity levels even when the trade-off is strongly specialist-favoring (community iv in Fig. 5A-C).

### Evolved communities use substitutable resources more efficiently than essential ones

Total biomass of evolved communities, and thus total consumer production, shows two major patterns: first, in any given landscape, evolved communities attain a higher total biomass when they feed on substitutable rather than essential resources; second, total biomass is positively related to the degree to which the affinity trade-off favors specialists, but only when resources are substitutable (Fig. 6A).

**Figure 6:**
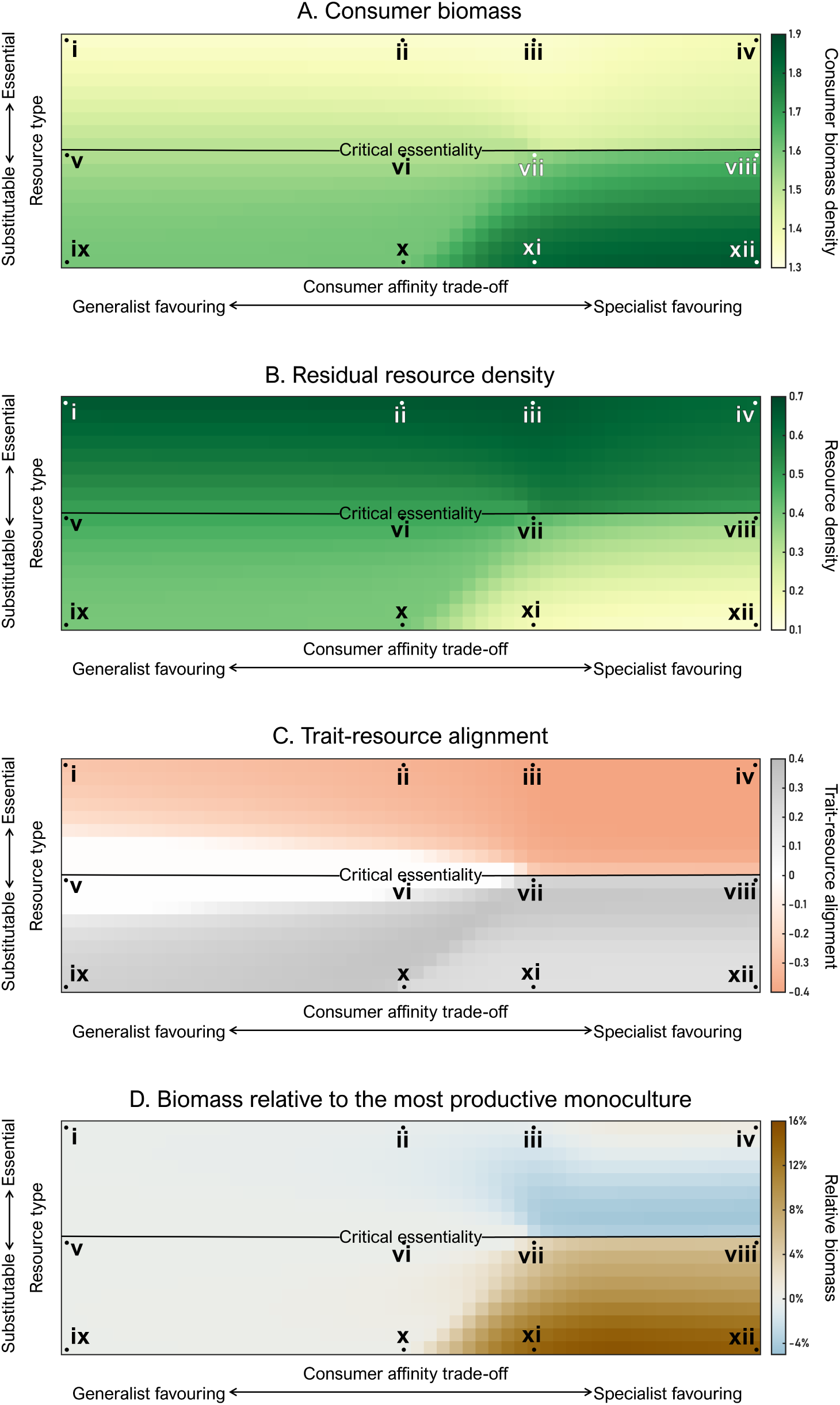
(A) Total biomass, (B) residual resource density, and (C) trait-resource alignment of the evolved consumer communities shown in Fig. 5 as functions of the strength of the affinity trade-off and the degree of resource substitutability. Also shown is (D) the biomass of the evolved communities relative to the most productive monocultures. Panels show mean values of four randomly generated resource supply landscapes with perfectly anti-correlated (ρ = −1) supplies of resources 1 and 2. The x- and y-axes scale as in Fig. 5A. Roman numerals indicate the locations of the histograms in Fig. 5C.

The first pattern is explained by the fundamental difference between consumers of essential vs. substitutable resources in how they align with the supply of the respective resource for which they have the higher affinity. Thus, because consumers of essential resources are locally competitively excluded from landscape patches with a high supply of the resource on which they are specialized (compare resource supply densities with specialist consumer densities in Fig. 4B), consumers of essential resources show negative trait-resource alignment (Fig. 6C). Consequently, when resources are essential, considerable amounts of both resources remain unused wherever they are in locally high supply (compare resource supply densities with residual resource densities in Fig. 4B), leading to low resource use efficiency and, thus, high residual resource densities and low consumer biomass (e.g. communities i-iv in Fig. 6A, B). In contrast, consumers of substitutable resources are positively aligned with the resource on which they are specialized (Fig. 6C; compare resource supply densities with specialist consumer densities in Fig. 4A), leading to high resource use efficiency, high consumer biomass (Fig. 6A), and low residual resource densities everywhere on the landscape (bottom panels in Fig. 4A; Fig. 6B).

The second pattern is explained by the broadening of the affinity trait space towards more strongly specialist-favoring trade-offs, which enables more specialized consumers of substitutable resources to transform an increasingly larger fraction of resources into consumer biomass. When resources are perfectly substitutable, a specialist-favoring trade-off drives evolution towards an endpoint with two extreme specialists (panels xi-xii in Fig. 5C), which, together, form the community with the highest possible resource use efficiency and, thus, generate both the highest biomass and lowest residual resource density under a given specialist-favoring trade-off (Fig. 6A, B). A similarly extreme specialization on a single resource is impossible when resources are essential (panels iii-iv in Fig. 5C), preventing consumers of essential resources from increasing their collective resource use efficiency and biomass even under the most strongly specialist-favoring trade-off (Fig. 6A, B).

### Adaptive evolution can cause transgressive overyielding or underyielding - depending on resource substitutability and trade-off strength

Comparing the consumer biomass of evolved communities in a given landscape against that of the corresponding monoculture with the highest biomass reveals, again, two patterns. First, when the trade-off is generalist-favoring, evolved communities perform similar to the best monoculture (polygon bounded by positions i-ii-x-ix in Fig. 6D). Second, when the trade-off is specialist-favoring, evolved communities can outperform the best monoculture considerably when resources are substitutable, but tend to underperform when resources are essential (polygon bounded by positions iii-iv-xii-xi in Fig. 6D).

The first pattern arises because, when the trade-off is generalist-favoring, the best monoculture is always the perfect generalist, regardless of resource substitutability. Evolved communities including (mild) specialists will therefore perform locally better in landscape patches where the resource supply is strongly skewed, but will underperform in landscape patches where the supply with the two resources is more balanced, the net effect across the entire landscape being an overall similar performance of evolved communities and the generalist monoculture.

When the trade-off is specialist-favoring, the best monoculture is, again, the perfect generalist when resources are substitutable, but a mild specialist when resources are essential. In the former case, the endpoint of adaptive evolution is an overyielding 2-species community of two strong and complementary resource specialists which, together, use and depress the two resources most efficiently and therefore always outperform the most productive generalist monoculture (polygon bounded by positions x-xii-viii-vii in Fig. 6B and D). In contrast, when resources are essential, evolution is driven by competition for the resources that are locally in lowest supply, preventing less specialized mutants with potentially higher resource use efficiency from being included in the community. Consequently, the best monoculture is a mild specialist, and the evolved community is often underyielding compared to it(blue areas inside the polygon bounded by positions ii-iv-viii-vii in Fig. 6B and D).

### Spatial heterogeneity in resource supply reduces consumer biomass – but only when resources are essential

Comparing the consumer biomass of evolved com_1_m_9_unities in spatially heterogeneous landscapes with that of communities that evolved in the corresponding spatially homogeneous landscape (with identical average resource supply densities *K*_1_ = *K*_2_), reveals a substantial negative effect of spatial heterogeneity on consumer biomass when resources are essential but not when resources are substitutable (Fig. 7). Wickman et al. (2019) have shown that adaptive evolution in spatially homogeneous landscapes with resource supply densities *K*_1_ = *K*_2_ has three possible outcomes: (1) a monoculture of a perfect generalist when the affinity trade-off is generalist- to mildly specialist-favoring (regions 1a and 1b in Fig. 7), (2) a monoculture of a mild specialist when the trade-off is more strongly specialist-favoring and resources are essential (region 2), and (3) two complementary, strong specialists when the trade-off is specialist-favoring and resources are substitutable (region 3). It then follows from the negative trait-resource alignment observed above the critical essentiality boundary (Fig. 6C) that evolved communities of consumers of essential resources have lower resource use efficiency and biomass in heterogeneous landscapes compared to the monoculture that is best adapted to the singular niche in the corresponding homogeneous landscape. In contrast, positive trait-resource alignment below the critical essentiality (Fig. 6C) enables evolved communities of consumers of substitutable resources to exploit locally skewed resource supply densities efficiently and, thus, reach similar resource use efficiency and biomass in heterogeneous as in homogeneous landscapes.

**Figure 7:**
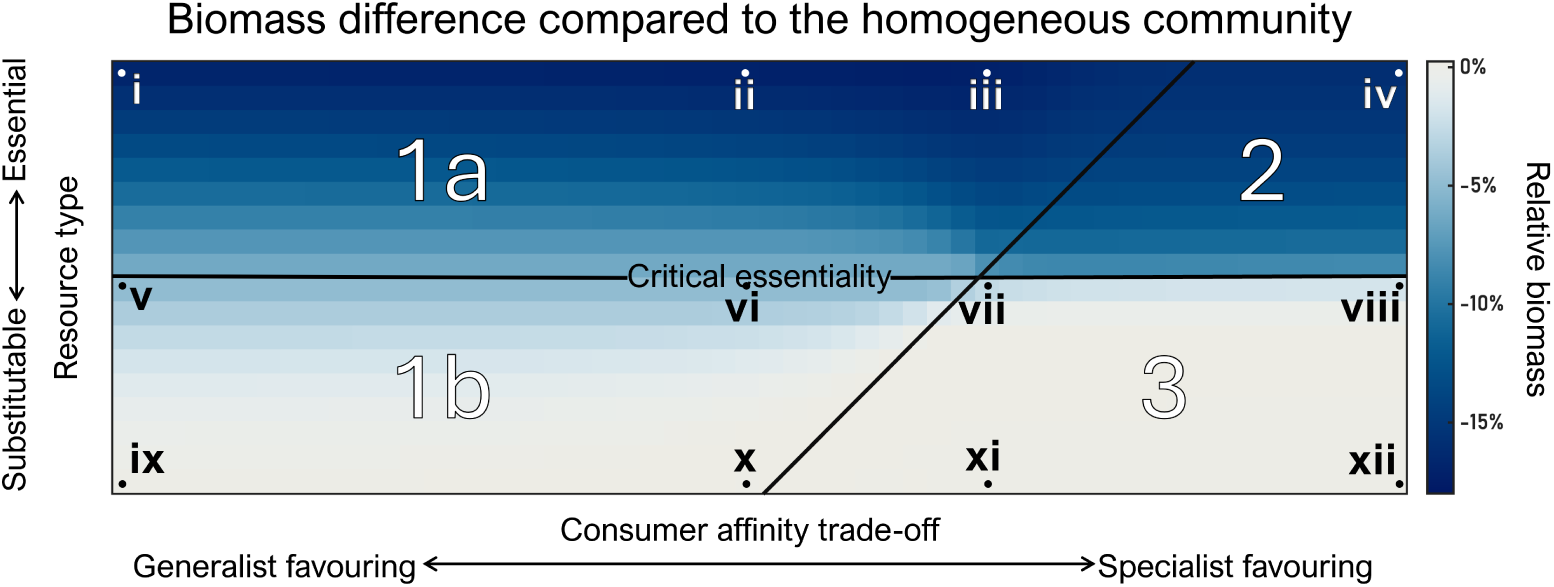
Percentage difference in biomass between evolved communities in spatially heterogeneous compared to homogeneous resource supply landscapes as a function of the strength of the consumer affinity trade-off and the degree of resource substitutability. Panels show mean values of the same four randomly generated resource supply landscapes with perfectly anti-correlated resource supplies yielding the results in Fig. 5 and 6. Arabic numbers separated by black lines indicate parameter regions yielding distinct evolutionary outcomes in the homogeneous supply landscape (see text). The x- and y-axes scale as in Fig. 5A. Roman numerals indicate the locations of the histograms in Fig. 5C.

### Effects of resource substitutability and trade-off shape are weaker in landscapes where resource supply densities are positively correlated

While the interactive effects of resource substitutability and trade-off shape on consumer diversity, biomass and resource use efficiency are qualitatively similar across all investigated resource supply landscapes, the patterns are strongest in landscapes where the supplies with resources 1 and 2 are negatively correlated, slightly weaker where the supplies are uncorrelated, but much weaker in landscapes where the supplies are strongly positively correlated (Fig. 8; for details see figs. 5-7 and figs. C.1-C.6). Specifically, as the spatial correlation between resource supplies increases, consumer biomass and resource use efficiency remain essentially unchanged in communities of consumers of substitutable resources but change considerably in communities of consumers of essential resources (compare histograms above and below the horizontal line indicating critical essentiality in Fig. 8C, D). However, the effect of correlation between resource supplies on species richness is seen in all but substitutable resources and in sufficiently specialist-favoring trade-off regions (compare the top and left histograms with the bottom-right one in Fig. 8A).

**Figure 8:**
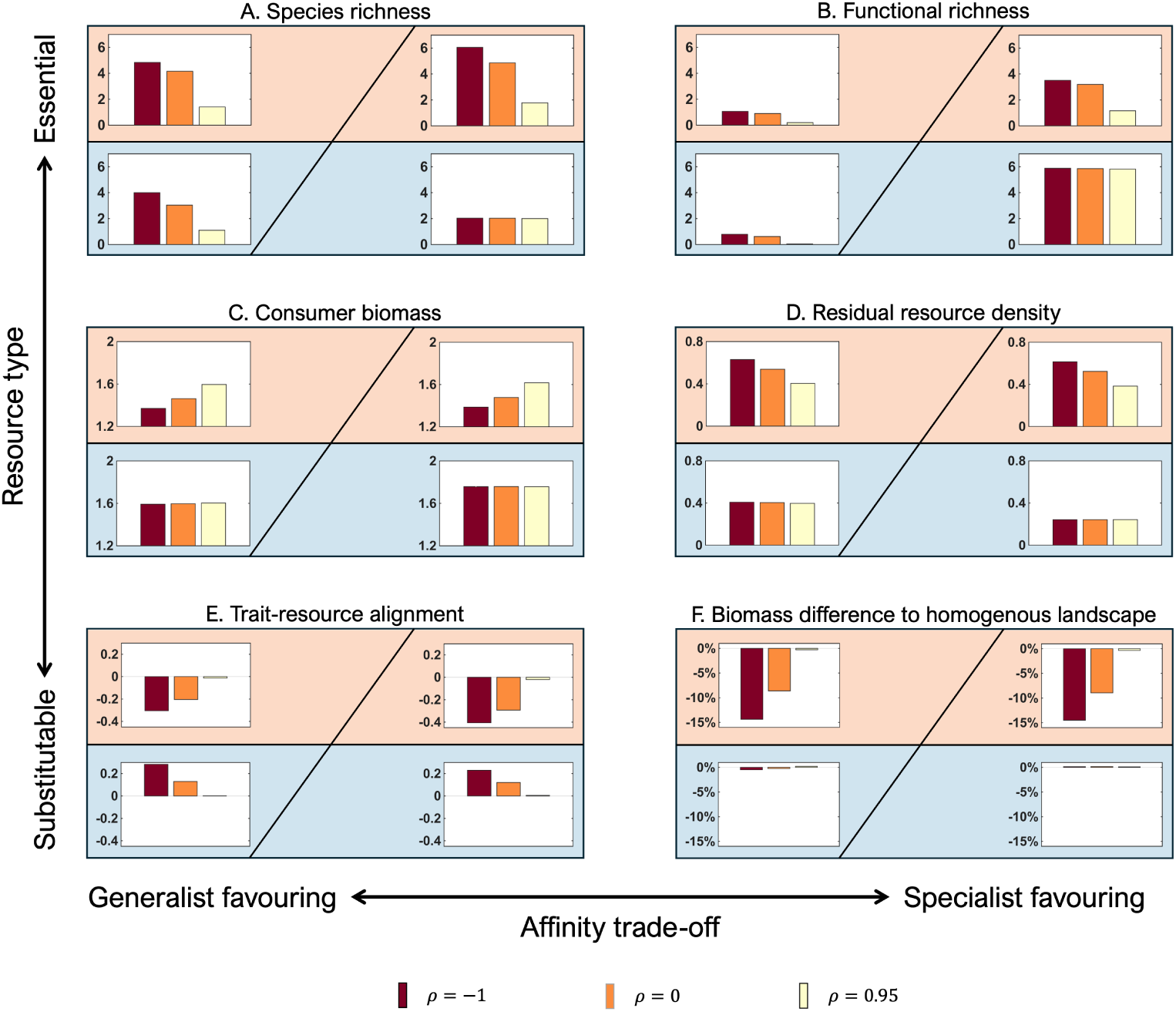
Summary of the effects of the landscape-wide cross-correlation between the local supplies with resources 1 and 2 on (A) species richness, (B) functional richness, (C) consumer biomass, (D) residual resource density, (E) trait-resource alignment, and (F) the biomass difference compared to a spatially homogeneous landscape. Bars show average values across the four regions in resource type - trade-off space indicated in Fig. 7, where the horizontal broken line corresponds to the critical essentiality separating essential resources (red background) from substitutable resources (blue background). Bar colors indicate landscapes with perfect negative cross-correlation (red, ρ = −1), no correlation (orange, ρ = 0), and strong positive correlation (cream, ρ = 0.95) between the local supplies of the two resources.

The underlying mechanism is as follows. Greater correlation between resource supplies reduces the prevalence of niches with strongly imbalanced resource supply ratios. This, in turn, reduces the degree of specialization among consumers of essential resources, such that evolved communities become progressively less specialized (Fig. 8B) and less species-rich (Fig. 8A) the more strongly positively correlated the supply ratios of the two resources are in the landscape. This occurs because highly specialized consumers of essential resources can only persist in landscapes including habitats with strongly imbalanced resource supplies; in the absence of such imbalances, these specialists cannot evolve. Since resource use efficiency for essential resources is most compromised in strongly imbalanced supply niches, as reflected in the negative trait-resource alignment above the critical essentiality (Fig. 8E), the removal of these imbalanced niches raises overall resource use efficiency (Fig. 8D) and, in turn, biomass (Fig. 8C) in landscapes where resource supplies are positively correlated. In contrast, owing to the positive trait-resource alignment below the critical essentiality (Fig. 8E), communities of consumers of substitutable resources already exploit imbalanced niches efficiently; a reduction in such niches therefore has little effect on consumer biomass and resource use efficiency (Fig. 8C, D). Finally, a strong positive correlation of the supplies with resources 1 and 2 implies that spatial heterogeneity almost exclusively reflects variation in absolute resource supply densities, while the supply ratios are near-identical and close to 1 everywhere on the landscape. Spatial heterogeneity therefore ceases to have a negative effect on consumer biomass when the correlation coefficient of resource supplies approaches 1 (Fig. 8F).

## 4 Discussion

In a first attempt towards the development of an evolutionary productivity-diversity theory, we explored how species richness, functional richness, total biomass, and resource use efficiency of evolved metacommunities of resource competitors depend on the degree of resource substitutability and the shape of the affinity trade-off. We found that, across this two-dimensional parameter space, conditions that promote the evolution of taxonomically diverse communities do typically not engender higher functional diversity, community biomass, and/or resource use efficiency. Thus, the highest community biomass, resource use efficiency and functional diversity emerge when resources are (near-) perfectly substitutable and the affinity trade-off is strongly specialist-favoring, which selects for a species-poor community of the two most extreme specialists. In contrast, the highest species diversity emerges when resources are (interactively) essential, and the affinity trade-off is mildly specialist-favoring, which selects for clusters of mild to intermediate specialists on each resource, yielding lower biomass and resource use efficiency – especially in landscape patches with imbalanced resource supply ratios.

### Evolutionary responses to substitutable versus essential resources diverge at the critical essentiality

The above results suggest that, starting from a single generalist phenotype, the evolution of species richness and community biomass will proceed in different – typically opposite – directions depending on whether consumers compete for substitutable vs. essential resources. This is consistent with the discovery of the ‘critical essentiality’, i.e., the boundary between substitutable-like and essential-like resources by Wickman et al. (2019). In heterogeneous landscapes, this boundary acts as a bifurcation point between opposite directionalities of local selection. Thus, when resources are substitutable, consumers diversify to occupy regions where the resource for which they have the higher affinity is in dominant supply, whereas the opposite is true when resources are essential. We found that this dichotomy in the community assembly pattern has significant consequences for all investigated response variables, including biomass and resource use efficiency (both being lower for essential than for substitutable resources), the trait-resource alignment (being negative for essential resources and positive for substitutable ones), and the impact of spatial heterogeneity on biomass (being negative for essential resources and near-neutral for substitutable ones). Moreover, the critical essentiality boundary and its immediate vicinity are themselves poor in biodiversity, allowing for a maximum of only two phenotypes when the trade-off is sufficiently specialist-favoring, and for no diversification at all when the trade-off is generalist-favoring.

Given the fundamental influence of the critical essentiality on the direction of local selection, the evaluation of biodiversity-ecosystem function relationships in competitive communities requires a good understanding of the degree of substitutability of the resources for which the consumers compete. Determining whether resources are above or below the critical essentiality will frequently require elaborate data collection, in particular for heterotrophic consumers which often feed on multiple biotic resources containing vital nutritional compounds (such as essential elements, amino acids, polyunsaturated fatty acids or vitamins) in greatly differing proportions (Sterner et al. 2002; Anderson et al. 2004; Raubenheimer et al. 2004; Sperfeld et al. 2012).

### Species richness does not equal functional richness

A point of contention in the biodiversity-ecosystem functioning debate has been to what extent ecosystem functioning is dependent on the species richness of a given community vs. its functional richness, i.e., the functional trait diversity of its constituent species (Hillebrand et al. 2009). A relevant step in addressing this question is to explore how well species richness and functional trait diversity are related. In our study, where consumer affinity was the trait under selection, species richness of the evolved communities turned out to be a poor predictor of their functional richness. The possibility of such a decoupling arises from the definition of functional richness as a convex hull, which, in a single trait dimension, reduces functional richness to the range spanned by the most extreme phenotypes present regardless of how many species occupy that range (Mason et al. 2005; Mouillot et al. 2005) or Rao’s entropy, which calculates the average trait dissimilarity between two randomly chosen individuals from the community (Rao 1982; Botta-Dukát 2005). As a result, a community with few species but extreme specialist phenotypes at opposite ends of the trait axis can have higher functional richness than a more species-rich community composed of more similar, less specialized phenotypes.

Our results contrast with the often positive relationships between species richness and functional richness reported from biodiversity manipulations, in which functional richness is not an emergent property of community assembly but a designed feature of the experiment (Naeem 2002; Cadotte et al. 2011). In our model, by contrast, functional richness emerges from evolutionary dynamics under a given trade-off structure and need not scale with the number of coexisting species at all. Our findings suggest that no single relationship between species richness and functional richness applies universally across communities (Díaz et al. 2001; Cadotte et al. 2011). Depending on the trade-off structure and resource substitutability characterizing a given lineage, diversification can proceed either through the emergence of few, functionally divergent specialists or through the proliferation of many functionally similar species.

### Productivity correlates with functional richness - but only when resources are substitutable

If species richness and functional richness are not universally coupled, it is natural to ask whether functional richness itself reliably predicts community productivity, or whether this relationship is similarly contingent on context. Functional richness has been reported to predict consumer biomass better than species richness in several studies (Tilman et al. 1997; Mokany et al. 2008), and niche complementarity is the mechanism most commonly invoked to explain why functionally diverse communities tend to be more productive (Díaz et al. 2001; Striebel et al. 2009; Wang et al. 2021; Amyntas et al. 2023). Our results suggest that the validity of this relationship depends on whether resources are substitutable or essential. Thus, when resources are substitutable, a specialist’s performance advantage is realized in those landscape patches where its preferred resource is in relatively higher supply. This generates a positive trait-resource alignment and, thus, niche complementarity, enabling different specialists to dominate in different regions of the landscape where each of them is more productive than a generalist would be in their place. The result is the evolution of a community that becomes simultaneously more functionally specialized and more productive. In contrast, when resources are essential, consumer growth remains constrained by whichever resource is locally scarcest. Specialization on a locally abundant resource then does not confer any fitness advantage, and selection drives specialization for the resource that is locally in shortest supply, generating negative trait-resource alignment. The result is a community that becomes more functionally specialized without becoming more productive.

### When does evolution lead to over-vs. underyielding

Our results paint a complex picture of evolved diversity-productivity relationships. We found that evolved metacommunities can be more, less, or as productive as the most productive monoculture, with this relationship depending strongly on both the consumer affinity trade-off and resource substitutability. This shows that evolution need not lead to transgressive overyielding, consistent with prior work on evolution (Matsuda et al. 1994; Metz et al. 2008; Rohr et al. 2023). Thus, while relative productivity differences between evolved metacommunities and the most productive monoculture are minimal under generalist-favoring trade-offs, specialist-favoring trade-offs can over- or underyield. Specifically, when resources are substitutable, local coexistence of extreme specialists allows evolved metacommunities to outperform the most productive monoculture locally everywhere in the landscape. Remarkably, these overyielding communities consist of only two species. Conversely, when resources are essential, evolved metacommunities do frequently underyield, yet underyielding is very modest (*≤* 4%). In line with experimental observations (Cardinale et al. 2007; Striebel et al. 2009), our analyses thus suggest that (empirically detectable) underyielding is an unlikely outcome also when communities are assembled by adaptive evolution. More at odds with some of the same studies, they also highlight that while overyielding is a possible evolutionary outcome, evolved species richness is counterintuitively low in such cases.

### The productivity cost of spatial heterogeneity

We explored the consequences of spatial heterogeneity on the relationship between consumer diversity and productivity in two distinct ways: first, by comparing spatially heterogeneous with spatially homogeneous landscapes; and second, by varying the sign and magnitude of the cross-correlation between the two resource supplies across the spatially heterogeneous landscapes. These analyses revealed that the effects of spatial heterogeneity differ qualitatively between substitutable and essential resources. Thus, when resources are substitutable, spatial heterogeneity has a moderate positive influence on species richness but no influence on biomass. These patterns arise because, when resources are substitutable and the landscape-averaged resource supply is balanced, evolution selects for similar phenotypes regardless of the degree of spatial heterogeneity, i.e., one to few similar-performing (near-)generalists when the trade-off is generalist-favoring, or the same two extreme specialists when the trade-off is specialist-favoring. In contrast, when resources are essential, both aspects of spatial heterogeneity can be strongly positively related to species richness and negatively to biomass, indicating a productivity cost of spatial heterogeneity. This productivity cost is highest in landscapes where the resource supplies are negatively correlated but negligible in landscapes where the supplies are strongly positively correlated. These patterns arise because trait-resource alignment is negative, and the resulting resource use efficiency is strongly reduced in landscapes including patches with greatly imbalanced resource supply ratios (Ptacnik et al. 2010; Hodapp et al. 2016), but not in landscapes where the resource supply ratio is balanced everywhere.

Overall, these findings suggest that the impacts of spatial heterogeneity on the diversity and productivity of consumers should be largest when resources are essential and their supplies are negatively correlated in space, making it relevant to ascertain the degrees of substitutability and spatial cross-correlation of different resources in empirical biodiversity-ecosystem functioning studies. For example, essential mineral nutrient supply is always negatively correlated with photosynthetically active radiation along pelagic depth gradients, which are often characterized by vertical niche partitioning of diverse phytoplankton communities (Latasa et al. 2017; Lofton et al. 2020). In line with the model expectation that such systems should be underyielding, artificial vertical mixing was shown to reduce phytoplankton taxonomic diversity while increasing overall community biomass (Jäger et al. 2008; Grossart et al. 2026). In contrast, the essential plant nutrients nitrogen and phosphorus tend to be positively correlated in space in both aquatic and terrestrial environments (Downing et al. 1992; Liu et al. 2013). In agreement with our model expectations, these systems become both more productive and more species poor typically when they are enriched with nitrogen and phosphorus such that the nutrient supply ratio becomes more balanced (Proulx et al. 1996; Seabloom et al. 2021).

### Conclusion and outlook

We have explored the role of adaptive evolution in generating biodiversity in metacommunities of resource competitors and the resulting relationships between evolved biodiversity and ecosystem functioning. In striking contrast to conventional wisdom and experiments with randomly assembled communities, but in agreement with observational studies and models of metacommunity dynamics (see references in the introduction), we found that, across the two-parameter space explored in this study, conditions that promote the evolution of diverse communities frequently do not engender higher functional richness, consumer biomass, or resource use efficiency. This suggests that the fundamental drivers of these measures differ or affect them differently, and we identified the critical essentiality as a key phenomenon separating parameter regions with fundamentally different relationships between biodiversity and ecosystem functions. Thus, when resources are (interactively) essential, both generalist- and specialist-favoring trade-offs select for phenotypes that prevent efficient resource partitioning locally, leading to negative trait-resource alignment and, consequently, low resource use efficiency and consumer biomass – both of which are independent of species richness (which is driven by the interplay of species sorting and mass effects) and functional richness (which is driven by the affinity trade-off). In contrast, when resources are complementary or fully substitutable, a more specialist-favoring affinity trade-off selects for phenotypes that efficiently partition resources locally, promoting a metacommunity of only two species characterized by high functional diversity, resource use efficiency, and biomass.

Introducing an evolutionary metacommunity perspective to classical biodiversity-ecosystem functioning research opens several further avenues for exploration. For example, we have treated resource substitutability in an entirely phenomenological way. Given the overarching importance of the concept of critical essentiality in our analyses, a more mechanistic approach is desirable to make this concept more easily applicable to empirical data. A next step in that direction could be to redefine resource substitutability in our model in terms of the stoichiometry of each resource’s content of essential nutritional compounds (Sterner et al. 2002; Sperfeld et al. 2012). Similarly, we only scratched the surface of exploring how spatial heterogeneity in resource supply affects evolved biodiversity, consumer biomass, and resource use efficiency. While we explored the full range of spatial cross-correlation patterns in resource supply, we kept other spatial dimensions constant. Future work should more systematically explore additional dimensions such as overall variance, directional trends and spatial autocorrelation. Finally, we focused on biodiversity and ecosystem functions at the spatial scale of entire metacommunities. Yet, the spatial scales of most biodiversity-ecosystem functioning studies are much smaller. A relevant extension of our model analyses would therefore be to explore how evolved biodiversity is related to ecosystem functions at the patch scale within different types of heterogeneous landscapes. Given the apparent mismatch between biodiversity-productivity relationships in experimental vs. observational studies, we believe that an evolutionary perspective can bring us closer to a better understanding of the biodiversity-productivity relationships found in mature ecosystems.

