## Supplementary for "Contrasting relationships between evolved consumer diversity and productivity for essential vs. substitutable resources"

### Appendix

#### Analytical derivation of the relationship between consumer biomass and resource densities

Let $\Omega = \mathbb{R}^2/\mathbb{Z}^2$ be the periodic two-dimensional torus. The area of the torus is normalized to be unity i.e., $\int \Omega \; dx = 1$. The ecological dynamics of the spatially explicit consumer-resource model on said torus is given by the following equations:

$$\frac{\partial A_i}{\partial t} = G_i A_i - \mu A_i + d\Delta A_i, \tag{S1a}$$

$$\frac{\partial R_j}{\partial t} = r_j \left(K_j - R_j\right) - \sum_{i=1}^{N} C_{ij} G_i A_i + d_R \Delta R_j, \quad j \in \{1, 2\}. \tag{S1b}$$

Without loss of generality, we assume that the biomass of the consumers and densities of the resources are unitless or share the same units and consider the total density as the sum of all the densities and biomass, i.e., $\mathcal{T} = \sum_{i=1}^{N} \int_\Omega A_i \; dx + \int_\Omega R_1 \; dx + \int_\Omega R_2 \; dx$. The rate of change of the total biomass is equal to the sum of the rates of change of densities and biomass,

$$\frac{\mathrm{d}\mathcal{T}}{\mathrm{d}t} = \sum_{i=1}^{N} \frac{\mathrm{d}A_i}{dt} + \frac{\mathrm{d}R_1}{dt} + \frac{\mathrm{d}R_2}{dt}. \tag{S2}$$

The right-hand side of eqn.S2 can be expanded as,

$$\frac{\mathrm{d}\mathcal{T}}{\mathrm{d}t} = \sum_{i=1}^{N} \frac{\mathrm{d}}{\mathrm{d}t} \left( \int_\Omega A_i \; \mathrm{d}x \right) + \frac{\mathrm{d}}{\mathrm{d}t} \left( \int_\Omega R_1 \; \mathrm{d}x \right) + \frac{\mathrm{d}}{\mathrm{d}t} \left( \int_\Omega R_2 \; \mathrm{d}x \right). \tag{S3}$$

Since the domain is fixed in time, we can use the Leibniz rule to differentiate under the integral. Thus, eqn.S3 transforms into

$$\frac{\mathrm{d}\mathcal{T}}{\mathrm{d}t} = \sum_{i=1}^{N} \int_\Omega \frac{\partial A_i}{\partial t} \; \mathrm{d}x + \int_\Omega \frac{\partial R_1}{\partial t} \; \mathrm{d}x + \int_\Omega \frac{\partial R_2}{\partial t} \; \mathrm{d}x. \tag{S4}$$

Substituting the equations for the rates of change of biomass and densities,

$$\frac{\mathrm{d}\mathcal{T}}{\mathrm{d}t} = \sum_{i=1}^{N} \int_\Omega \left(G_i A_i - \mu A_i + d\Delta A_i\right) \; \mathrm{d}x + \sum_{j=1}^{2} \int_\Omega \left( r_j \left(K_j - R_j\right) - \sum_{i=1}^{N} C_{ij} G_i A_i + d_R \Delta R_j \right) \; \mathrm{d}x. \tag{S5}$$

Since the boundary is periodic, the rates of change of diffusion cancel out over the surface by the divergence theorem. We also assume the ecological dynamics is at a dynamic equilibrium, resulting in the following simplification of eqn.S5,

$$0 = \sum_{i=1}^{N} \int_{\Omega} \left(G_i A_i - \mu A_i\right)\ \mathrm{d}x + \sum_{j=1}^{2} \int_{\Omega} \left( r_j \left(K_j - R_j\right) - \sum_{i=1}^{N} C_{ij} G_i A_i \right)\ \mathrm{d}x. \tag{S6}$$

Rearranging eqn.S6 results in,

$$\sum_{i=1}^{N} \int_{\Omega} \left( G_i A_i - \sum_{j=1}^{2} C_{ij} G_i A_i \right)\ \mathrm{d}x + \sum_{j=1}^{2} r_j \int_{\Omega} K_j\ \mathrm{d}x = \sum_{i=1}^{N} \mu \int_{\Omega} A_i\ \mathrm{d}x + \sum_{j=1}^{2} r_j \int_{\Omega} R_j\ \mathrm{d}x. \tag{S7}$$

The following simplifying assumptions result in the relationship between consumer biomass and resource densities seen in figures 6, S2 and S5:

1. Consumed resources are fully converted into consumer biomass, $C_{i1} + C_{i2} = 1$.

2. The renewal rates are independent of the resources and are unity, $r_j = 1$.

3. The mortality rates are independent of the consumer species and are unity $\mu = 1$.

4. The average resource supply of each resource is unity, $\int_{\Omega} K_j = 1$.

More explicitly, the above assumptions transform eqn.S7 to

$$2 = \sum_{i=1}^{N} \int_{\Omega} A_i\ \mathrm{d}x + \sum_{j=1}^{2} \int_{\Omega} R_j\ \mathrm{d}x. \tag{S8}$$

#### Numerical experiments and parameters

The spatially explicit consumer-resource model that is used for the ecological dynamics is given by eqn.1 and the evolutionary dynamics is achieved by solving eqn.8 together with eqn.1. To increase computational efficiency, the numerical implementation of the ecological and evolutionary dynamics was done using a scaled version of the equations. This scaling allowed us to eliminate the dimensions of all parameters and thus reduce the number of parameters by the number of unique dimensions.

The scaled spatially explicit consumer-resource model is given by the following equation,

$$\frac{\partial \hat{A}_i}{\partial \hat{t}} = \hat{G}_i \hat{A}_i - \hat{\mu} \hat{A}_i + \hat{d} \Delta \hat{A}_i, \tag{S9a}$$

$$\frac{\partial \hat{R}_j}{\partial \hat{t}} = \hat{r}_j \left( \hat{K}_j - \hat{R}_j \right) - \sum_{i=1}^{N} C_{ij} \hat{G}_i \hat{A}_i + \hat{d}_R \Delta \hat{R}_j, \quad j \in \{1, 2\}, \tag{S9b}$$

where the scaled per-capita growth function is

$$\hat{G}_i = \hat{G}_{\max} \frac{\left(\left(\hat{a}_1(\chi_i)\hat{R}_1\right)^s + \left(\hat{a}_2(\chi_i)\hat{R}_2\right)^s\right)^{\frac{1}{s}}}{2^{\frac{1}{s}}\hat{H} + \left(\left(\hat{a}_1(\chi_i)\hat{R}_1\right)^s + \left(\hat{a}_2(\chi_i)\hat{R}_2\right)^s\right)^{\frac{1}{s}}}. \tag{S10}$$

***Table S1:*** *Scaled variables used for the numerical simulations, their modification from their original equations and their values or ranges. Here,K is the average resource supply over the entire landscape and L is the length of one side of the "boundary" of the landscape.*

| New variable | Equation | Value |
|---|---|---|
| $\hat{A}_i$ | $A_i \cdot K^{-1}$ | – |
| $\hat{R}_j$ | $R_j \cdot K^{-1}$ | – |
| $\hat{K}_j$ | $K_j \cdot K^{-1}$ | – |
| $\hat{t}$ | $t \cdot (G_{\max} - \mu)$ | – |
| $\hat{G}_i$ | $G_i \cdot (G_{\max} - \mu)^{-1}$ | – |
| $\hat{G}_{\max}$ | $G_{\max} \cdot (G_{\max} - \mu)^{-1}$ | 2 |
| $\hat{H}$ | $G_{\max} \cdot a_{\text{norm}}^{-1} \cdot K^{-1}$ | 0.2 |
| $\hat{\mu}$ | $\hat{G}_{\max} - 1$ | 1 |
| $\hat{r}_j$ | $r_j \cdot (G_{\max} - \mu)^{-1}$ | 1 |
| $\hat{d}$ | $d \cdot (G_{\max} - \mu)^{-1} \cdot L^{-2}$ | $1 \cdot 10^{-4}$ |
| $\hat{d}_R$ | $d_R \cdot (G_{\max} - \mu)^{-1} \cdot L^{-2}$ | $1 \cdot 10^{-4}$ |
| $\hat{a}_j$ | $a_j \cdot a_{\text{norm}}^{-1}$ | – |
| $a_0$ | – | $[-5.95, 24.23]$ |
| $b$ | – | 4 |
| $\chi_i$ | – | $[-1, 1]$ |
| $s$ | – | $[-1, 1]$ |

We discretized the space into a $32 \times 32$ grid and used the method of lines *i.e,* converting the partial differential equations into a system of coupled ordinary differential equations, to solve the spatially explicit system. We used a pseudo-spectral method (Trefethen 2000) to calculate the resultant diffusion matrix, the discrete version of the Laplacian or diffusion function $\Delta$. We solved for the ecological steady state by simultaneously solving $1024 \times (n + 2)$ coupled ODE equations (where $n$ is the number of consumer species) using the Backward Differentiation Formula of the CVODE solver in the SUNDIALS package in Julia 1.9.3.

We simulated the evolutionary dynamics of the system by simultaneous solving $1024 \times (n+2)$ coupled ODE equations dictating the ecological dynamics and $n$ equations dictating the trait evolution using the KenCarp4 solver in the DifferentialEquations package in Julia 1.9.3. To model the trait evolution, we dynamically calculated the selection gradient using the perturbation method proposed by Wickman et al. (2017).

We used MATLAB2025a to compile all the results and produce all the figures.

#### Results for additional supply landscapes

This appendix provides figures and a description of the results obtained for landscapes in which the supplies of resources 1 and 2 are uncorrelated (Fig. S1-S3) and strongly positively correlated (Fig. S4-S6) in space in the same format as the corresponding results for anti-correlated resource supplies presented in the main text (Fig. 5-7).

Overall, the patterns in species richness and consumer biomass are qualitatively similar across all three degrees of spatial cross-correlation, but their magnitudes are correlation-dependent. Compared to anti-correlated landscapes (Fig. 5A), species richness is slightly lower in uncorrelated landscapes (Fig. S1A) and substantially lower in strongly correlated landscapes (Fig. S4A), suggesting a generally decreasing relationship between the degree of resource cross-correlation and species richness (see also Fig. 8A). Functional richness shows a similarly decreasing trend with increasing degree of resource supply correlation, except in the region of parameter space dominated by extreme specialists (the bottom-right area in Figs. 5B, S1B and S4B), which is unaffected by changes in cross-correlation, confirming that spatial heterogeneity does not affect community composition in that part of parameter space. In contrast, consumer biomass and resource use efficiency of evolved communities increase with increasing cross-correlation, especially in environments with essential resources (figs. 6A-B, S2A-B, and S5A-B).

The patterns in trait-resource alignment, as well as the comparisons of evolved metacommunity biomass with both the most productive monoculture and the corresponding community that evolved in a spatially homogeneous environment are also qualitatively similar across all three degrees of spatial cross-correlation; with the exception of the biomass comparison with the most productive monoculture in the region of parameter space dominated by extreme specialists, the patterns weaken, again, gradually towards environments with increasingly positive cross-correlation of the resource supplies to effectively disappear near the endpoint of perfect cross-correlation (figs. 6C-D, 7, S2C-D, S3, S5C-D, S6; see also Fig. 8E-F).

Taken together, these results indicate that the main qualitative conclusions from our model analyses on how the species richness of adaptively evolved metacommunities relates to functional diversity, resource use efficiency, and community biomass hold regardless of the correlation structure between resource landscapes, though the magnitude of these effects depends systematically on the degree of cross-correlation.

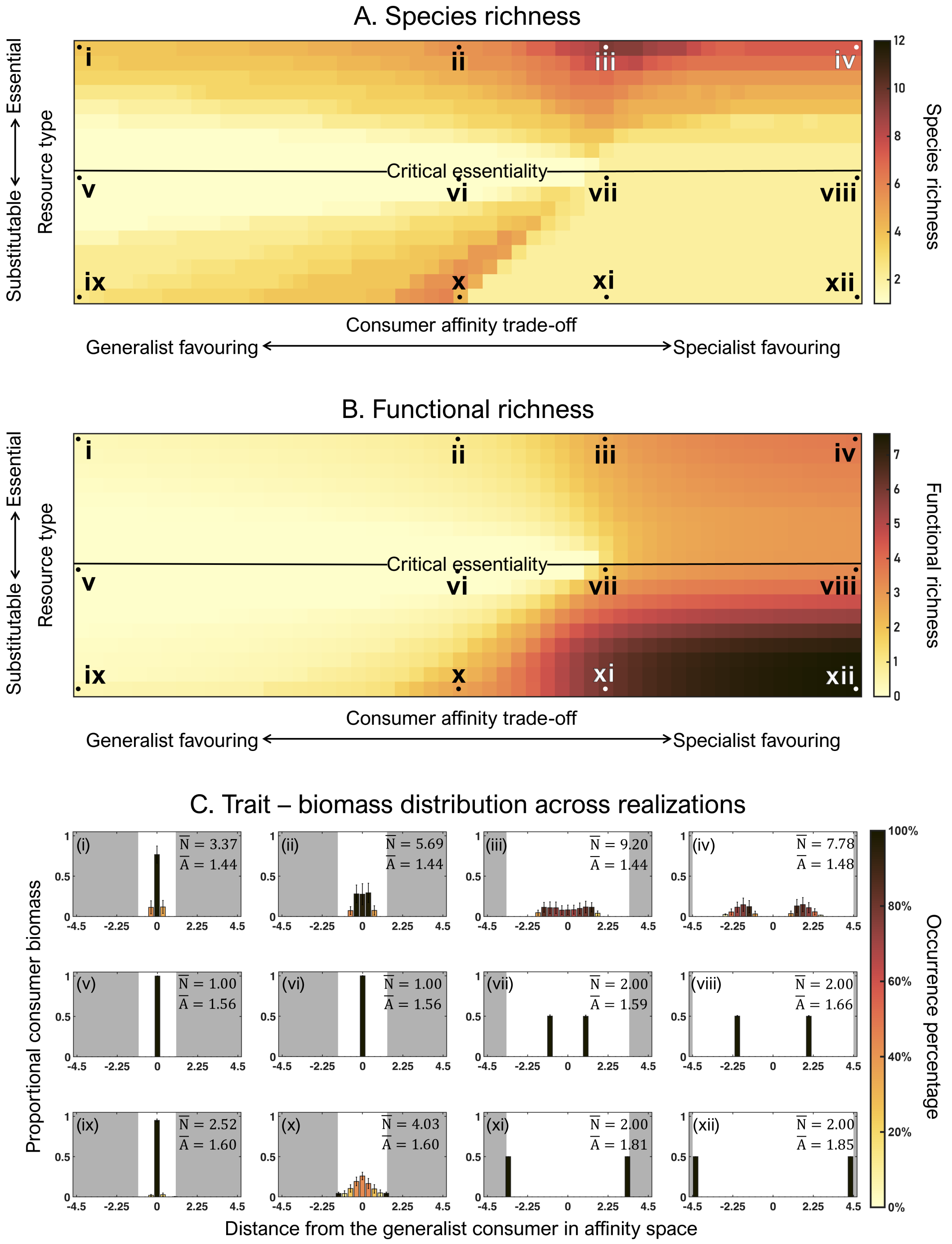


***Figure S1:*** *Evolved (A) species richness, and (B) functional richness and (C) trait distribution in landscapes with uncorrelated ($\rho = 0$) supplies of resources 1 and 2 and for different combinations of the strength of the affinity trade-off (ranging from strongly generalist- to strongly specialist-favouring, see fig. 2A) and the degree of resource substitutability (ranging from perfectly substitutable to interactively essential, see Fig. 1A and B). Panels A and B show mean values of four randomly generated resource supply landscapes, with the central horizontal black lines indicating the critical resource essentiality. Histograms in panel C show the distribution of consumer biomass in affinity trait space of evolved metacommunities across 160 randomly generated landscapes for the 12 combinations of affinity trade-off strength and degree of resource substitutability indicated by Roman numerals in panels A and B. S and U indicate the average values of species richness and total consumer biomass across all 160 realizations. Columns represent the average proportional biomass (+1 standard deviation) of consumers that are specialists on resource 1 (negative x-axis values) and resource 2 (positive x-axis values), respectively, where the histograms' x-axes represent the specialists' distance in affinity space from the generalist consumer (with affinities $a_1 = a_2 = 1$), with higher absolute values indicating stronger specialization, and the bar color indicating the frequency of occurrence of an affinity distance across all 160 realizations. The white space around each histogram indicates the available trait space, where borders to the grey areas mark the maximum affinity values on a given trade-off curve.*

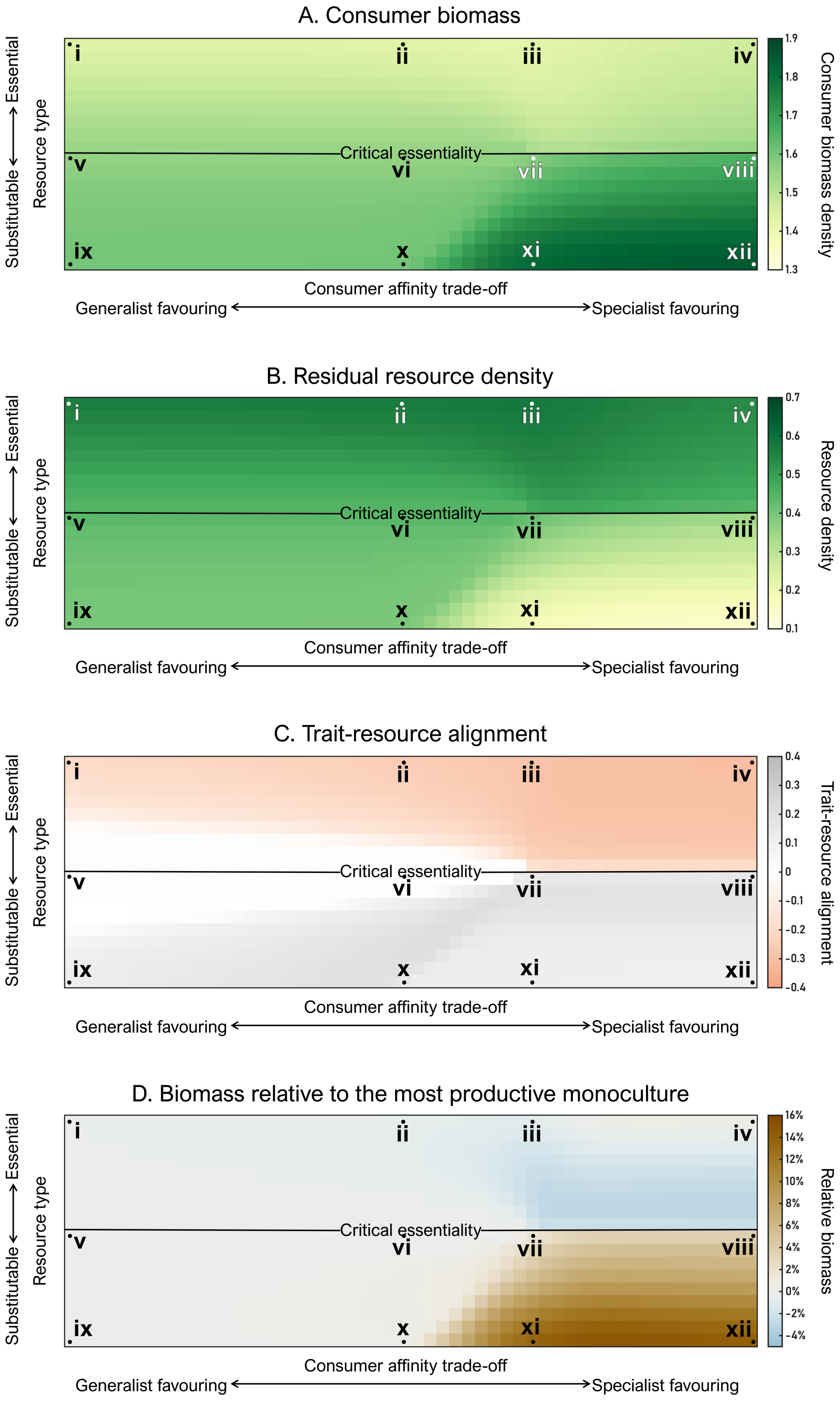


***Figure S2:*** *(A) Total biomass, (B) residual resource density, and (C) trait–resource alignment of the evolved consumer communities shown in Fig. S1 as functions of the strength of the affinity trade-off and the degree of resource substitutability. Also shown is (D) the biomass of the evolved communities relative to the most productive monocultures. Panels show mean values of four randomly generated resource supply landscapes with uncorrelated ($\rho = 0$) supplies of resources 1 and 2. The x- and y-axes scale as in Fig. S1A. Roman numerals indicate the locations of the histograms in Fig. S1C.*

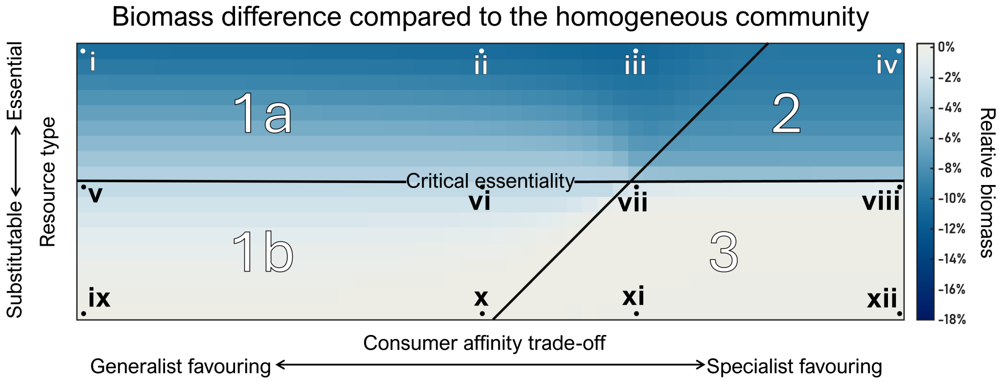


***Figure S3:*** *Percentage difference in biomass between evolved communities in spatially heterogeneous compared to homogeneous resource supply landscapes as a function of the strength of the consumer affinity trade-off and the degree of resource substitutability. Panels show mean values of the same four randomly generated resource supply landscapes with uncorrelated resource supplies yielding the results in Fig. S1 and S2. Arabic numbers separated by black lines indicate parameter regions yielding distinct evolutionary outcomes in the homogeneous supply landscape (see text). The x- and y-axes scale as in Fig. S1A. Roman numerals indicate the locations of the histograms in Fig. S1C.*

Wickman, Jonas, Sebastian Diehl, Bernd Blasius, et al. 2017. "Determining Selection across Heterogeneous Landscapes: A Perturbation-Based Method and Its Application to Modeling Evolution in Space". *The American Naturalist* 189 (4): 381–395. https://doi.org/10.1086/690908.

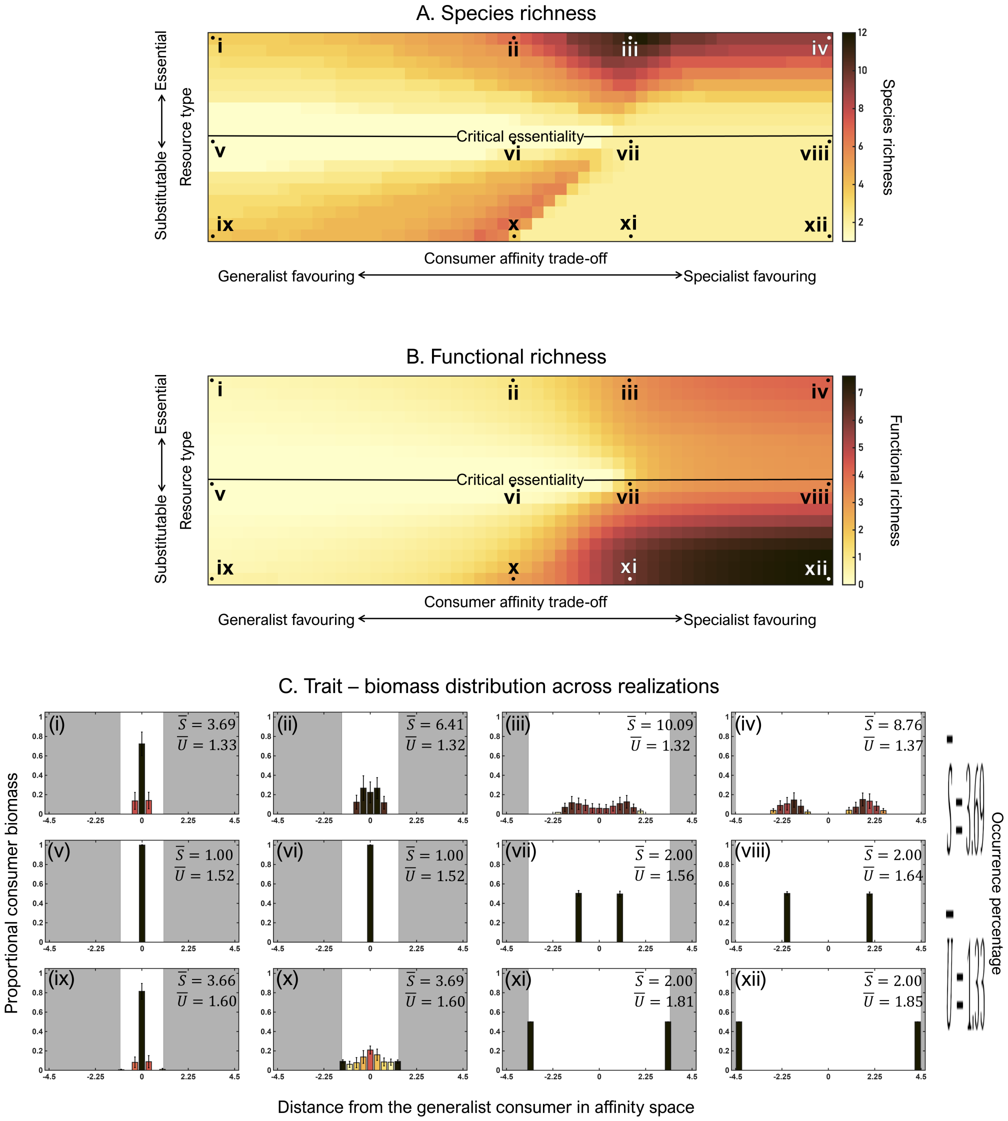


***Figure S4:*** *Evolved (A) species richness, and (B) functional richness and (C) trait distribution in landscapes with highly correlated($\rho = 0.95$) supplies of resources 1 and 2 and for different combinations of the strength of the affinity trade-off (ranging from strongly generalist- to strongly specialist-favouring, see fig. 2A) and the degree of resource substitutability (ranging from perfectly substitutable to interactively essential, see Fig. 1A and B). Panels A and B show mean values of four randomly generated resource supply landscapes, with the central horizontal black lines indicating the critical resource essentiality. Histograms in panel C show the distribution of consumer biomass in affinity trait space of evolved metacommunities across 160 randomly generated landscapes for the 12 combinations of affinity trade-off strength and degree of resource substitutability indicated by Roman numerals in panels A and B. S and U indicate the average values of species richness and total consumer biomass across all 160 realizations. Columns represent the average proportional biomass (+1 standard deviation) of consumers that are specialists on resource 1 (negative x-axis values) and resource 2 (positive x-axis values), respectively, where the histograms' x-axes represent the specialists' distance in affinity space from the generalist consumer (with affinities $a_1 = a_2 = 1$), with higher absolute values indicating stronger specialization, and the bar color indicating the frequency of occurrence of an affinity distance across all 160 realizations. The white space around each histogram indicates the available trait space, where borders to the grey areas mark the maximum affinity values on a given trade-off curve.*

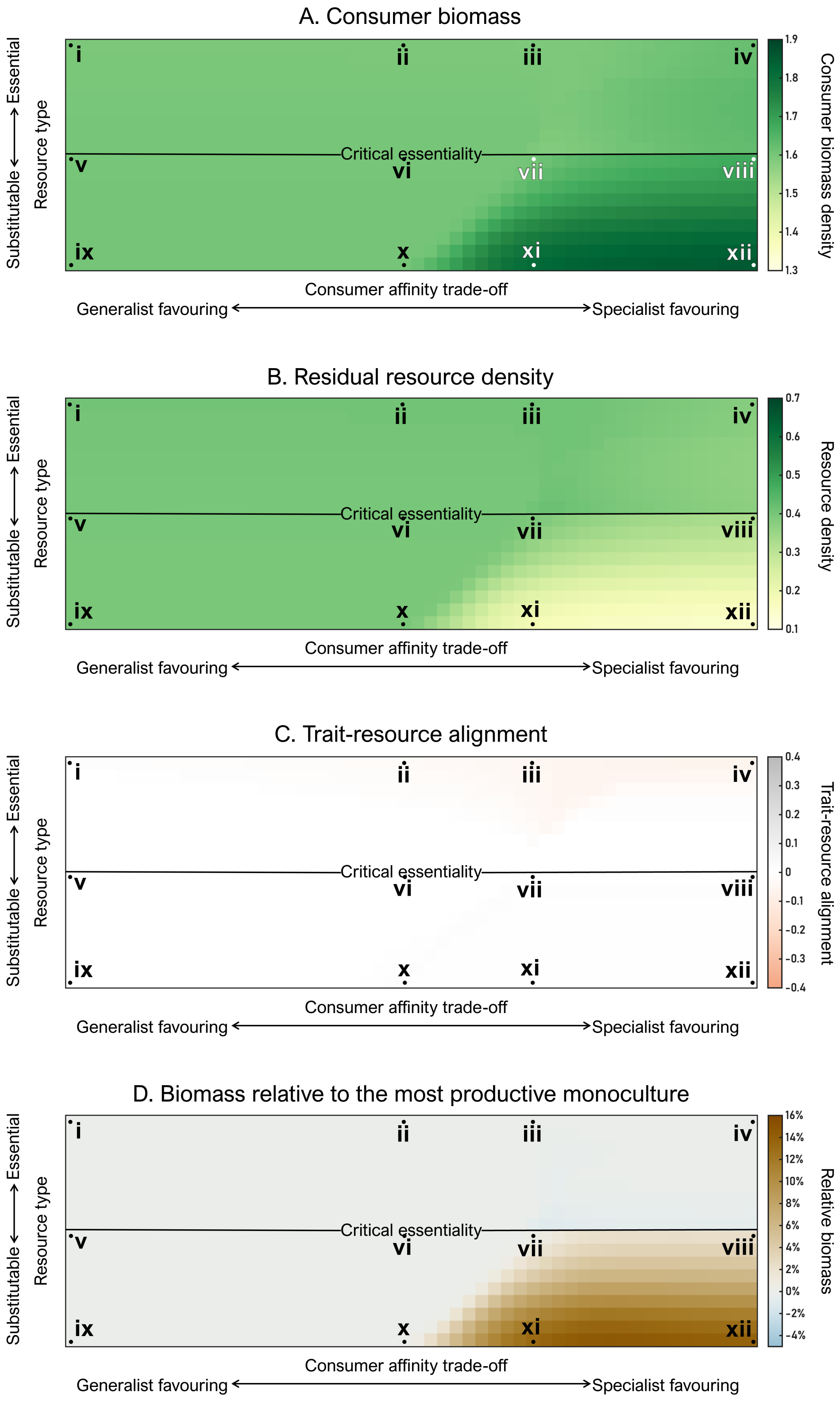


***Figure S5:*** *(A) Total biomass, (B) residual resource density, and (C) trait–resource alignment of the evolved consumer communities shown in Fig. S4 as functions of the strength of the affinity trade-off and the degree of resource substitutability. Also shown is (D) the biomass of the evolved communities relative to the most productive monocultures. Panels show mean values of four randomly generated resource supply landscapes with highly correlated ($\rho = 0.95$) supplies of resources 1 and 2. The $x$- and $y$-axes scale as in Fig. S4A. Roman numerals indicate the locations of the histograms in Fig. S4C.*

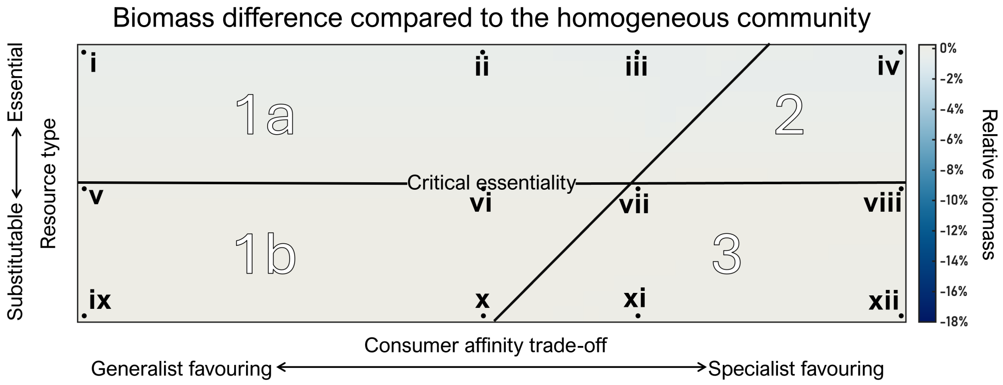


***Figure S6:*** *Percentage difference in biomass between evolved communities in spatially heterogeneous compared to homogeneous resource supply landscapes as a function of the strength of the consumer affinity trade-off and the degree of resource substitutability. Panels show mean values of the same four randomly generated resource supply landscapes with highly correlated resource supplies yielding the results in Fig. S4 and S5. Arabic numbers separated by black lines indicate parameter regions yielding distinct evolutionary outcomes in the homogeneous supply landscape (see text). The x- and y-axes scale as in Fig. S4A. Roman numerals indicate the locations of the histograms in Fig. S4C.*
